# Adipose tissue as a site of immune activation and dysfunction in individuals with obesity and asthma

**DOI:** 10.64898/2026.06.08.730463

**Authors:** Dawn C. Newcomb, Alessandra Tomasello, Jean-Philippe Cartailler, Bilgunay Ilkin Safa, LaToya Hannah, Shristi Shrestha, Shahar Hartman, Melissa H Bloodworth, Kevin Niswender, John R. Koethe, Samuel Bailin, James Matthew Luther, Nancy J Brown, Mona Mashayekhi, Katherine N Cahill

## Abstract

Obesity increases local inflammatory responses in adipose tissue. Individuals with obesity have increased asthma incidence and severity and reduced responses to asthma therapeutics through unknown mechanisms. To identify mechanisms by which increased fat mass augments asthma pathogenesis, single cell RNA sequencing of the immune-rich stromovascular fraction of subcutaneous adipose tissue was conducted from well-characterized adults with obesity-associated asthma matched to adults without asthma. Individuals with asthma had increased abundance of perivascular macrophages and lymphoid-associated macrophages (LAMs) and reduced abundance of classical monocytes and CD4+ and CD8+ naïve T cells. Pseudo-bulk differential expression (DE) identified upregulation of cellular metabolism, specifically oxidative phosphorylation, and decreased immune homeostatic pathways in asthma across immune cell subsets. Cell type specific DE analysis of effector cell subtypes identified significant induction of metallothionein gene expression in asthma, a signature of immune cell dysfunction characterized by both an activation and exhaustion phenotype. Gene co-expression analysis identified gene modules associated with asthma diagnosis, lung function, and biomarkers of type 2 inflammation were enriched in effector cells. These data identify adipose tissue dysfunction occurs in obesity-associated asthma and support adipose tissue as therapeutic target to address the enhanced asthma risk among those with obesity.

**Grant Support:** NIH U01AI155299, P30DK020593, R01AI182159, K23HL159351, UL1RR024975-03, P30CA68485, P30EY08126, G20RR030956, 5UL1TR002243, KL2TR002245, P30AI110527, DK020593, American Heart Association 17SFRN33520017.

## Introduction

Nearly half of adults are projected to be obese by 2030 (1). Increased obesity rates presage higher rates of obesity-associated complications, including asthma. Obesity is a risk factor for incident asthma across the life span and a risk factor for asthma severity, with a clear dose response relationship (2–6). Individuals with obesity-associated asthma have an increased risk of exacerbations requiring systemic corticosteroids and hospitalization and poorer symptom control. Compared to lean individuals with asthma, those with comorbid obesity, defined by elevated body mass index (BMI), demonstrate impaired response to inhaled corticosteroids and often fail to achieve adequate asthma control with cytokine-targeted biologic treatments (7–9). Obesity-related metabolic dysfunction including insulin resistance (IR), hyperglycemia, and systemic inflammation, marked by elevated interleukin (IL)-6 and CRP, are associated with asthma exacerbation risk and increased lung function decline over time (10, 11). Several lines of evidence show that adipose tissue (AT) dysfunction is a critical regulator of metabolic dysregulation and systemic inflammation (12). However, our understanding of the interplay between AT inflammation and asthma pathogenesis and asthma control is entirely unknown.

Type (T)2 inflammation, characterized by group 2 innate lymphoid cells (ILCs), T helper type 2 (Th2) cells and elevated IL-4, IL-5, and IL-13 cytokines, are thought to promote AT immune homeostasis in lean states (13–16). Increased adiposity results in an altered immune landscape in the AT with more inflammatory CD8+ T cells and lipid associated macrophages (LAMs) and decreased numbers of Th2 cells and ILC2 cells (17, 18). However, in asthma, T2 inflammation is strongly linked to airway inflammation, characterized by increased T2 cell subsets such as Th2, ILC2, and eosinophils and increased T2 cytokines including IL-4, IL-5, and IL-13. These asthma biomarkers can be detected in the airway and peripheral blood. In murine obesity models of asthma and clinical asthma with comorbid obesity, a dampening of classical T2 inflammatory processes is observed, characterized by decreased peripheral blood eosinophils and exhaled nitric oxide (FeNO), and an increase in neutrophilic, Th1 and Th17 signatures, particularly among females (19–23). Interestingly, T cell subsets in the peripheral blood correlate with T cell subsets in AT in some studies (24). Several pro-inflammatory factors linked with AT dysfunction, including IL-6, are associated with severity in asthma regardless of anthropometric measurements (10, 25) Interestingly, AT Th2 cell abundance has been shown to inversely correlate with IR, hsCRP, and IL-6 (24). An interplay between AT inflammation and the chronic airway lung inflammation from asthma is hypothesized given an overlap in inflammatory signatures but remains poorly understood.

Elucidating changes in AT immune cell composition, communication, and cytokine production in obesity-associated asthma may identify novel therapeutic targets to treat this increasingly prevalent and often refractory condition. We hypothesized that patients with asthma and comorbid obesity have a unique AT immune signature compared to patients with obesity without asthma. Characterizing the unique inflammatory environment of asthma in obesity is imperative to understanding the mechanistic relationship between obesity and increased asthma prevalence and severity, augmented airway inflammation, and reduced responses to currently available asthma therapeutics.

## Results

Stromovascular AT samples were chosen from cohorts of adults with obesity, with (n=16) and without (n=16) comorbid asthma, enrolled into prospective clinical trials at an academic medical center between 2017 and 2024. Samples were balanced on sex, race/ethnicity, BMI, hemoglobin A1C, hemostatic model of insulin resistance (HOMA-IR), and cardiovascular risk (Table 1). The participants with asthma had moderate-to-severe, persistent, uncontrolled disease defined by daily medium-or-high dose inhaled corticosteroid (ICS) use and an asthma control questionnaire (ACQ)-6 score ≥1.5 (Table 2) (26, 27).

**Table 1.** Cohort Demographics and clinical parameters.

| Demographics | Obese, No asthma | Obese, Asthma |
| --- | --- | --- |
|  | n=16 | n=16 |
| Age (yr) | 49.5 ± 14.1 | 50.0 ± 14.4 |
| Sex (female, no. [%]) | 12 (75) | 12 (75) |
| Race (White) | 11 (68.8) | 12 (75) |
| BMI (kg/m<sup>2</sup>) | 38.7 ± 6.1 | 39.0 ± 6.9 |
| A1c (%) | 5.8 ± 0.3 | 5.6 ± 0.4 |
| Total cholesterol (mg/dl) | 188.9 ± 28.10 | 190 ± 36.62 |
| HDL (mg/dl) | 47.5 ± 9.61 | 45.5 ± 9.23 |
| SBP (mmHg) | 125.1 ± 11.99 | 122.9 ± 9.45 |
| eGFR (mL/min/1.73 m<sup>2</sup>) | 87.68 ± 19.10 | 94.06 ± 13.85 |
| PREVENT score (% 10-year risk) | 4.4 ± 3.4 | 4.8 ± 4.0 |
| PREVENT score (% 30-year risk) | 20.6 ± 11.2 | 21.1 ± 12.1 |

**Table 2.** – Asthma cohort clinical features.

| **Asthma Clinical Characteristics** | **Mean ± SD** |
| --- | --- |
| Age at diagnosis (yr) | 32.7 ± 22.1 |
| Asthma duration (yr) | 17.3 ± 18.5 |
| FEV1 (% of predicted) | 69.9 ± 16.8 |
| FEV1/FVC Ratio | 0.70 ± 0.11 |
| ACQ-6 score | 2.5 ± 0.4 |
| AQLQ score | 3.8 ± 0.9 |
| Blood eosinophils (cells/μl) | 267 ± 185 |
| FeNO (ppb) | 24.1 ± 15.5 |
| Total serum IgE (IU) | 131.6 ± 92.7 |
| Medium-dose ICS (no [%]) | 12 (75%) |
| High-dose ICS (no [%]) | 4 (25%) |
| Fluticasone Propionate (mcg equivalent/day) | 445 ± 275 |
| LABA use (no [%]) | 13 (81.3%) |
| LAMA use (no [%]) | 1 (6.3%) |
| Montelukast use (no [%]) | 5 (31.3%) |

### Subcutaneous adipose tissue immune cell abundance

Single cell RNA sequencing (scRNA-seq) was conducted on the immune-rich stromovascular AT fraction, and the analysis showed the expected immune cell signatures including myeloid and lymphoid populations (Figure 1A-D). Sample processing was utilized to isolate immune cells and remove mature adipocytes in the sample and therefore few remained for sequencing (Figure 1B). The myeloid and T/NK cells were annotated using characteristic markers to define cell subsets (Figure 1C, D, Supplemental Figure 1)(28). scCODA analysis established credible differences in immune cell enrichment between the asthma and non-asthma samples (29). Classical monocytes were enriched in the non-asthma samples, while LAMs and perivascular macrophages were enriched in asthma samples (Figure 1E). Non-asthma samples also had increased naïve CD4+ and CD8+ T cells (Figure 1F), while asthma samples demonstrated a trend toward increased T effector memory (TEM) cell numbers that did not achieve the credible difference threshold. Together, these data show that asthma and non-asthma AT samples have differential immune cell composition in both the myeloid and T/NK compartments with increased AT immune effector cells present in the setting of asthma.

**Figure 1.**
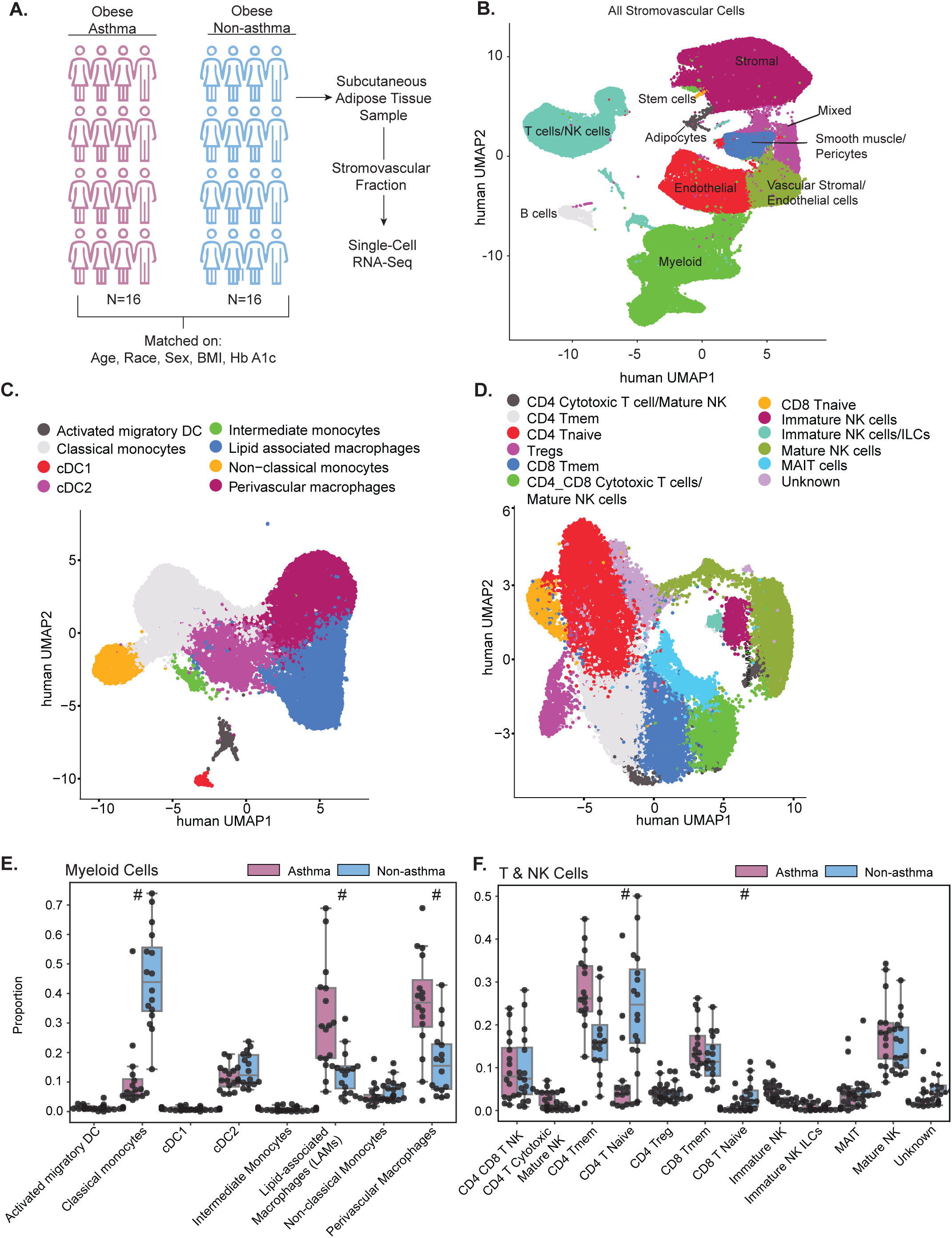
– Single cell landscape of non-asthma and asthma immune cell signatures in the stromal vascular fraction of subcutaneous adipose tissue identifies differential abundance of myeloid and T cell fraction based on asthma status in adults with obesity. (A) Study design and schematic of workflow for obtaining samples and running scRNA sequencing. (B) Global UMAP of cell types from non-asthma and asthma samples. (C-D) Annotated UMAP of myeloid (panel C) and T/NK cell subsets (panel D). (E-F) Relative abundance of myeloid and T/NK cell subsets by disease state determined by scCODA analysis. # represents credible differences between asthma and non-asthma groups via scCODA. DC – dendritic cell, ILC – innate lymphoid cell, NK – natural killer, TEM – T effector memory.

### AT gene signatures in asthma highlight the presence of activated and inflammatory CD4+ and CD8+ TEM cells

Differential gene expression analysis revealed that naïve CD8+ T cells from participants with asthma had modestly increased expression of *GZMK* and *CCL5/RANTES*, two genes associated with the promotion of inflammation in the respiratory tract (30). *LINC01871*, a marker of T cell activation, and *S100A4*, which is linked to dysfunctional effector function and increased proliferation capacity, were also increased in naïve CD8+ T cells from patients with asthma (Figure 2A) supporting an state of cellular activation(31). CD8 TEM demonstrated significant induction of *FABP4* and *XCL1*, two TEM survival signals (32, 33), the co-stimulatory receptor *TNFRSF9/4-1BB*, which is important in T cell expansion and cytokine production (34), *LINC01871*, and *BCL2A1*, a survival signal, in asthma (Figure 2B). In contrast, genes important in regulating differentiation and senescence (ID1) were suppressed in asthma.

**Figure 2.**
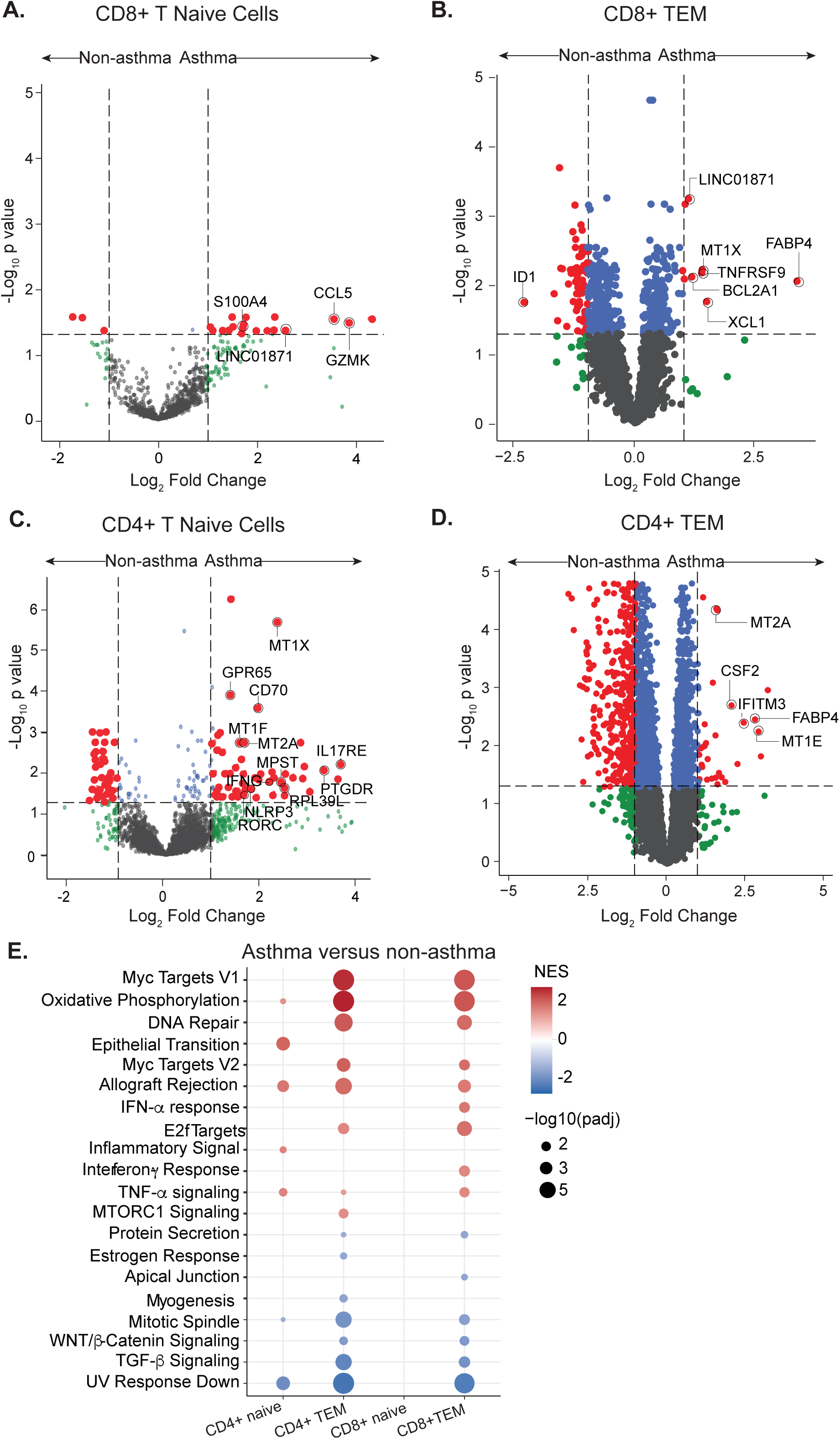
– Asthma AT expression signatures highlight that CD4+ and CD8+ cells are more activated and inflammatory in AT from obese patients with asthma compared to non-asthma obese controls. (A-D) Pseudo-bulk differential expression (DE) analysis of CD8+T naïve cells, CD8+ TEM cells, CD4+ T naïve cells, and CD4+ TEM cells with p-values adjusted for multiple testing and fold-change set at 1.0. (E) Merged Hallmark Gene set enrichment analysis (GSEA) of pseudo-bulk DE analysis with non-asthma set as the reference group across CD4+ and CD8+ naïve and TEMs. NES – normalized enrichment score.

Results from gene set enrichment analysis (GSEA) in CD8+ TEM demonstrated enrichment of MYC targets, oxidative phosphorylation, DNA repair, IFNα and IFNγ, and TNFα signaling via NFκB, and downregulation of TGFβ and WNT/β-Catenin pathways in AT from patients with asthma (Figure 2E). Recent work in severe asthma identified altered circulating and airway CD8+ T cells which are proliferative (35), including circulating CD8+ T cytotoxic cells with the ability to synthesize T2 cytokines (36). The AT CD8+ T cell signature supports extension of increased CD8+ T cell activation and increased effector function described in the circulation and airway to the AT in the setting of obesity-associated asthma.

Differential gene expression in naïve CD4+ T cells revealed the induction of genes associated with Th2 and Th17 polarization, both implicated in asthma pathogenesis, with Th17 cells implicated in females with asthma and obese-associated asthma (19, 22, 23). These genes include: *IL17RE*, a component of the IL-17 receptor; *PTGDR*, prostaglandin D receptor 2 or CRTH2; *NLRP3*, a transcriptional regulator of Th2 differentiation (37); *RORC* the master transcription factor for Th17, ILC3, and γδT cells (38); and *GPR65*, a promotor of Th1 and Th17 cells (39) (Figure 2C). Increased expression of the costimulatory marker *CD70* support the hypothesis that naïve CD4+ T cells are activated in the setting of asthma. Consistent with this, MsigDB C7 immunologic signature GSEA of naïve CD4+ T cells revealed the positive enrichment of CD4+ TEM and negative enrichment of naïve and central memory pathways (data not shown). Moreover genes which regulate mitochondrial function such as *MPST* (3-mercaptopyruvate sulfurtransferase) (40), and mitochondrial activity such as *RPL39L* (41), are induced in naïve CD4+ T cells from patients with asthma. In CD4+ TEM cells from patients with asthma, there is an upregulation of *IFITM3*, a regulator of Th2 cell differentiation (42), and *CSF2/GM-CSF*, which suggests impaired Treg cell suppression via IL-3 (43) (Figure 2D). Additionally, these DE results in CD4+ TEM cells from patients with asthma were associated with enrichment of oxidative phosphorylation, MYC targets, DNA repair and depletion of TGFβ and WNT/β-Catenin pathways on GSEA, mirroring what we observed in the CD8+ T cells and supporting increased CD4+ T cell activation with skewing toward T2/T17 phenotypes (Figure 2E).

Both CD4+ and CD8+ TEM cells from patients with asthma are characterized by *FABP4*, critical for FFA uptake, mitochondrial oxidative metabolism, and cell survival (44), and metallothionein (MTs) gene expression (Figure 2B, 2D). Normalized counts show that *MT1X* and *MT2A* are significantly increased in patients with asthma (Supplemental Figure 2A). MTs carry zinc and other heavy metals and have roles in heavy metal toxicity, DNA damage, and oxidative stress. Prior studies have shown that increased MT gene signatures are linked with T cell dysfunction characterized by dual activation and exhaustion signatures and are also increased in obese patients with type 2 diabetes compared to non-diabetic controls (45, 12). These genes may be of particular interest in the context of asthma, as their regulatory elements contain a glucocorticoid receptor-binding site. Prior studies in HeLa cell lines established dexamethasone induces MT gene expression and the induction of MT2A specifically disrupted the ability of Zn to inhibit NFκB activity (46, 47).

### Myeloid signatures in asthma highlight a heightened inflammatory state

The classical M1-M2 macrophage dichotomy often used to describe macrophage populations at sites of inflammation in the respiratory tract does not align with the current understanding of macrophage heterogeneity in adipose tissue, particularly in the obese state. While we observed that classical monocytes were depleted in asthma (Figure 1C, E), they showed the greatest number of upregulated genes as compared to non-asthma samples (Figure 3A). *HLA-DRA/B1/DPA1/DQA2* expression, markers of monocyte activation and antigen presentation capability which respond to cytokines IFNγ, GM-CSF, and IL4, and *APOC1/2* and *NRP2*, two transcripts upregulated during the differentiation process of a monocyte to a macrophage, showed high levels of induction in asthma. *LTC4S* and *FCER1a*, two transcripts with classical roles in T2 inflammation, allergy, and asthma, were also upregulated on classical monocytes in asthma samples and have previously been shown to be induced on monocytes by IL-13 and circulating IgE levels, respectively (48). In addition, *CXCL5*, a mediator of insulin resistance in AT and viral-induced chemotactic factor for neutrophils in the airway (49, 50), *CXCL9*, a chemotaxis factor for cytotoxic T/NK/macs with a reported role in T1 airway inflammation in asthma(51), and *IL7R*, a cytokine receptor induce by LPS and TNF exposure (52), are also induced in classical monocytes from participants with asthma. *CCL2/MCP-1*, a secreted cytokine with a role in monocyte recruitment to inflammatory sites, was also induced in asthma. GSEA revealed upregulation of pathways associated with stress (E2F targets and DNA repair), cell proliferation and differentiation (MYC targets), and metabolic pathways of oxidative phosphorylation in classical monocytes in asthma, while TGFβ, estrogen response, TNFα, WNT/β-catenin signaling, and apoptosis pathways were enriched in controls (Figure 3D). The KEGG asthma pathway was also enriched in classical monocytes from donors with asthma, further highlighting an inflammatory signature characteristic of asthma which is detectable in the AT (data not shown). These data support the reduction of classical monocytes in the AT may be the result of local differentiation to macrophages or a programming process in the adipose tissue which primes monocytes to contribute to non-T2 inflammatory processes at sites of inflammation outside of the adipose tissue.

**Figure 3.**
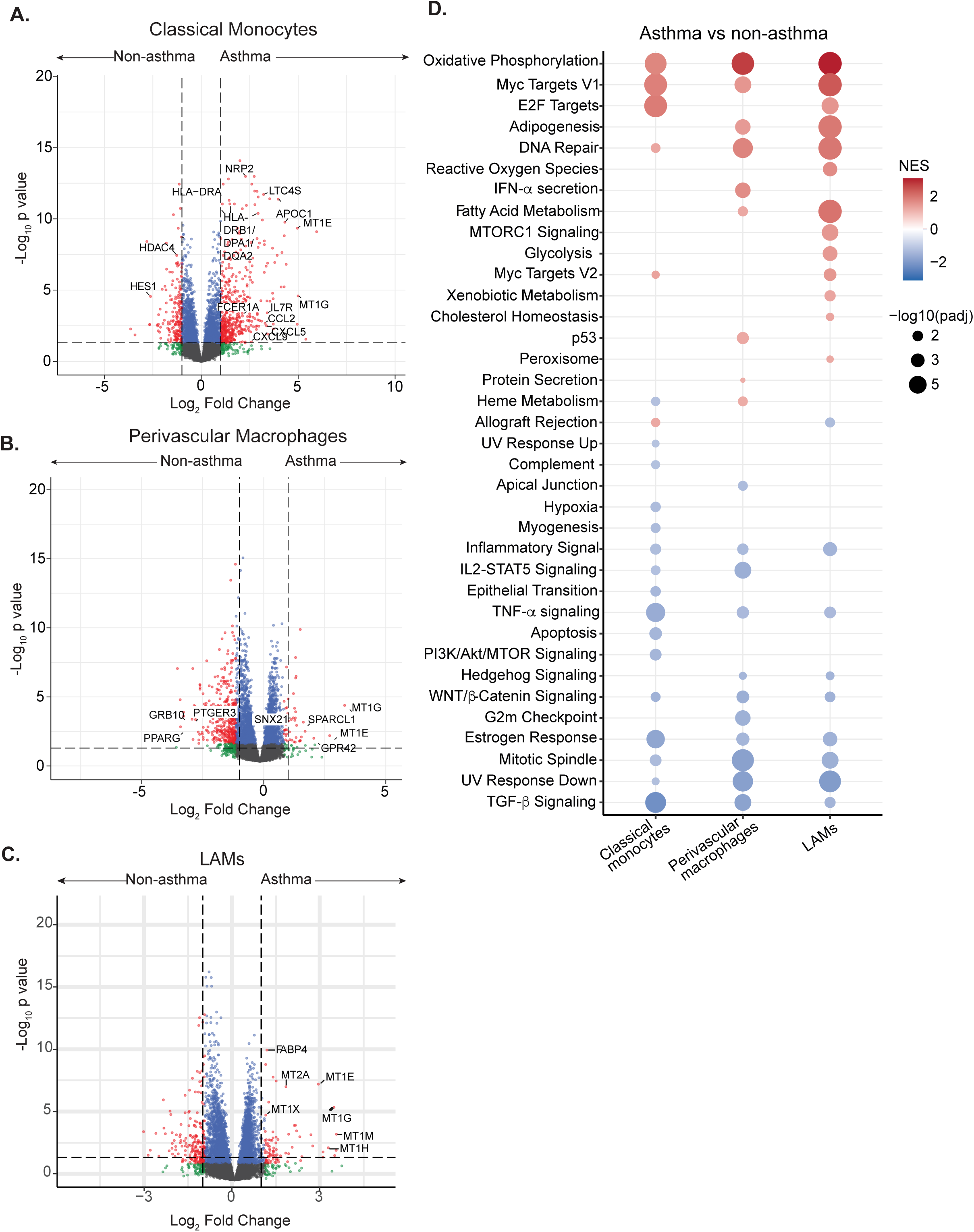
– Asthma AT genes in differentially abundant myeloid subsets show increased inflammatory signatures. (A-C) Pseudo-bulk differential expression (DE) analysis of classical monocytes, perivascular macrophages, and lipid-associated macrophages (LAMs) with p-values adjusted for multiple testing and fold-change set at 1.0. (D) Merged Hallmark Gene set enrichment analysis (GSEA) of pseudo-bulk DE analysis with non-asthma set as the reference group across differentially abundant myeloid subsets. NES – normal-ized enrichment score.

Perivascular macrophages (PVMs) are the dominant AT resident macrophage subgroup in lean AT (17, 18), yet intriguingly they are expanded in AT from participants with asthma (Figure 1E). PVM from asthma samples are characterized by downregulation of *GRB10*, a gene required for lipolysis and cold-induced thermogenesis (53), *PTGER3*, the loss of which is linked to high-fat diet-induced fat expansion (54), and *PPARG*, which is required for macrophage differentiation, polarization toward anti-inflammatory state, and lipid metabolism (Figure 3B). A modest upregulation of *GPR42/FFAR1*, the receptor for long-chain free fatty acids, and *SPARCL1*, a proinflammatory adipokine which is associated with non-alcoholic fatty liver disease, is observed (Figure 3B) (55). GSEA of the PVMs mirrored observations in the CD4/8+ TEM with enrichment of oxidative phosphorylation, p53, heme metabolism, and fatty acid metabolism pathways as well as pathways associated with stress, including DNA repair, and immune response, including IFNα signaling and MYC targets. PVM in asthma had reduced enrichment of the TGF-β, WNT/β, and IL2/STAT5 signaling pathways (Figure 3D). The functional significance of expanded PVMs in the obese state is not clear but may imply a macronutrient signature in the AT.

LAMs are enriched in AT during obesity-induced inflammation (56) and were expanded in AT from participants with asthma (Figure 1E). Similar to what was observed in the T cell compartment, MT genes were highly induced in the LAMs (Figure 3C and Supplemental Figure 2B). GSEA of LAMs demonstrated enrichment for metabolic pathways (oxidative phosphorylation, adipogenesis, fatty acid metabolism) and pathways associated with stress, including DNA repair, ROS pathways, and E2F targets in asthma (Figure 3D), like the PVMs. Enrichment of multiple immune response pathways in the non-asthma samples mirrored what was observed in the PVMs.

### Systemic biomarkers of inflammation inadequately capture AT inflammation in asthma

Adipose tissue is a source of various proinflammatory mediators which can be detected in systemic circulation. Interestingly, no difference was detected in serum high-sensitivity (hs)CRP or IL-6 levels. Systemic levels of two myeloid-associated inflammatory biomarkers, IL-1RA and MCP-1/CCL2 were significantly elevated or showed a trend toward elevation, respectively, in asthma. Systemic IL-8 levels were lower in the plasma of participants with asthma and GM-CSF and TNFα showed a similar trend. No difference was detected in plasma IL-18 and IL-1β levels, two markers along with IL-1rα of inflammasome activation, although levels for both mediators were very low. Similarly, IFNγ, IL-12p70, FGF21, and FABP4 levels did not differ by phenotype (Figure 4).

**Figure 4.**
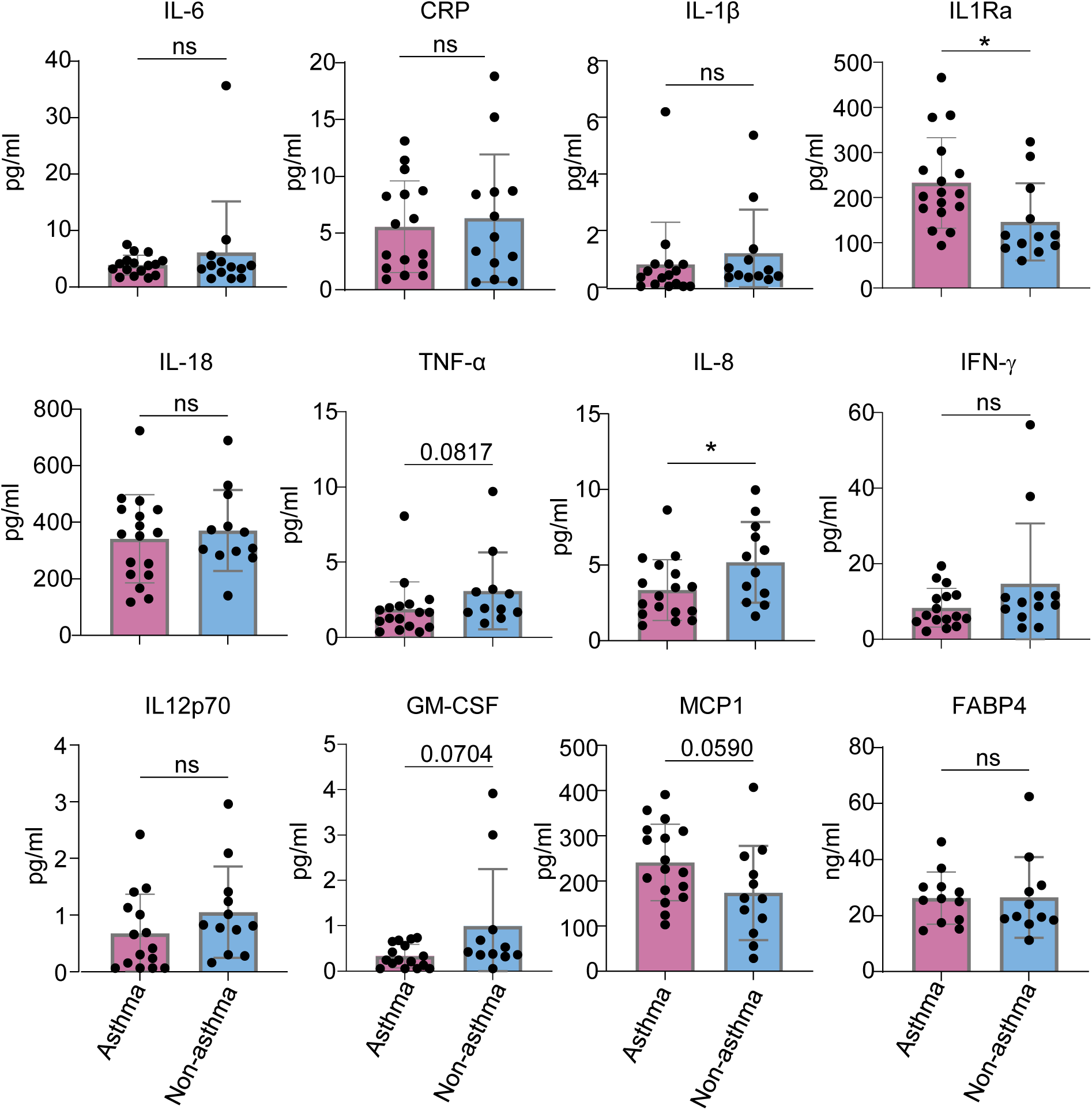
– Systemic biomarkers of inflammation inadequately capture the enhanced adipose tissue inflammation observed in asthma. Plasma cytokines are stratified by disease state. Wilcoxon test applied. * p<0.05, ns, non significant.

### T/NK Cell iterative WGCNA identifies asthma diagnosis as the key clinical factor driving gene expression in obese AT

Since patients with asthma had increased signatures of metabolism and stress-induced responses in many cell subsets within the myeloid and T/NK cell compartments, we were interested in determining if there was a clinical co-variable driving these observed differences. Iterative weighted gene co-expression network analysis (iterativeWGCNA) was conducted on T/NK cells from the total cohort (n=32) integrating clinical metabolic and asthma-specific metadata (Figure 5A-C). Co-expression modules were further classified into metamodules by community detection analysis. The main clinical variable driving gene expression correlation to form metamodules was asthma diagnosis, with the strongest correlation with modules in metamodule E (Figure 5A,B). Asthma surrogates such as bronchodilator induced changes in forced expiratory volume in 1 second (FEV1) positively correlated with metamodule E (purple color) and negatively correlated with metamodules C and D (green and red, respectively). Metamodule B (orange), particularly module B4, was correlated with biomarkers of T2 inflammation, blood and sputum eosinophils and serum periostin, but not asthma status. Module F20 was correlated with higher markers of systemic inflammation (total platelet count, triglyceride/HDL ratio, blood neutrophils), serum IgE value, and exacerbation frequency (number of oral corticosteroid courses in the prior year) but without correlation to asthma status. Interestingly, age, sex, race/ethnicity, BMI, metabolic markers of A1c, fasting insulin and glucose, fat distribution including visceral adipose tissue, cardiovascular disease risk (10- and 30-year PREVENT scores), and systemic biomarkers of inflammation demonstrated few or no correlations with gene expression modules.

**Figure 5.**
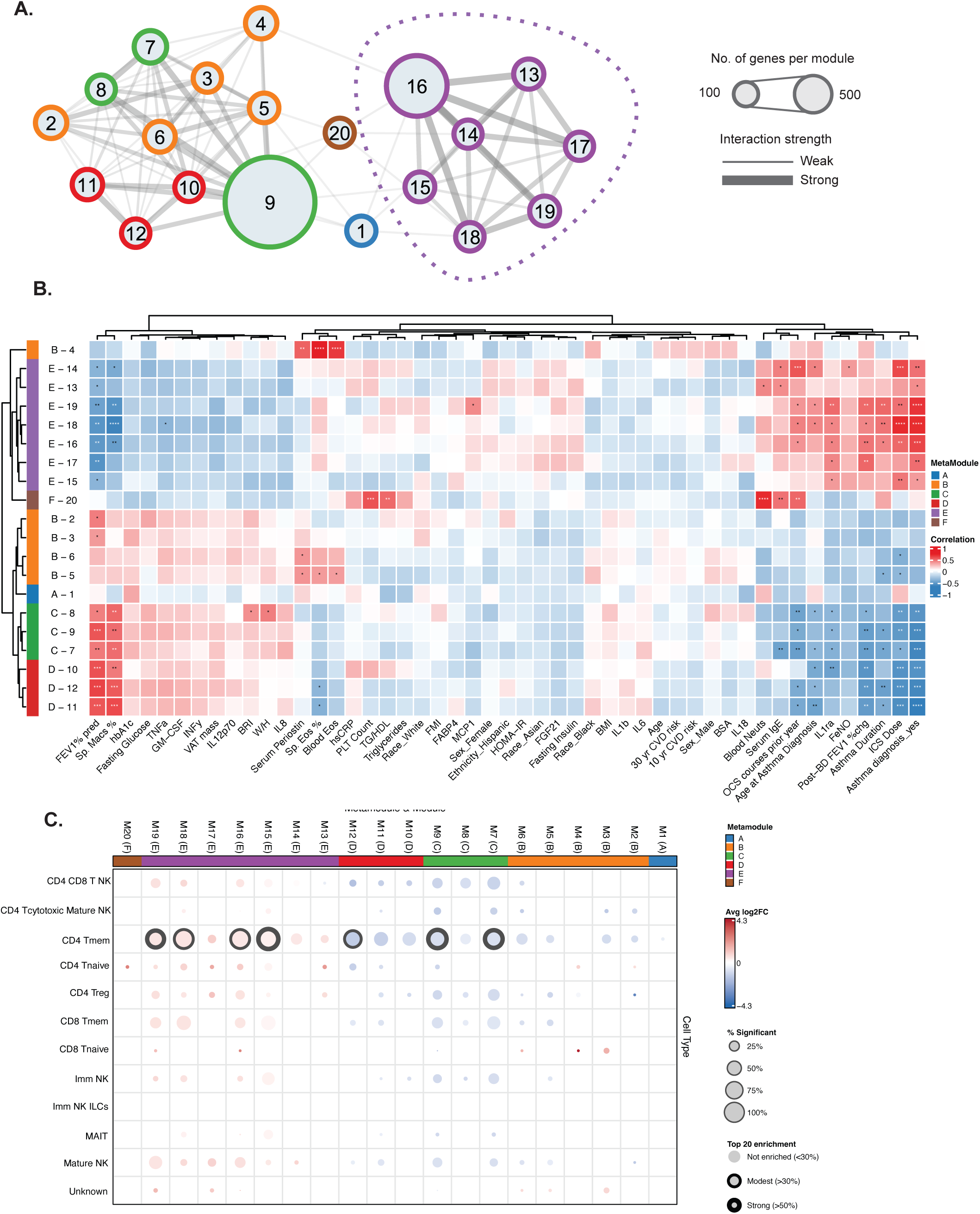
– Iterative Weighted gene co-expression analysis (iterativeWGCNA) of T/NK fraction identifies asthma as the clinical variable driving module formation. A) iterativeWGCNA module and metamodule network. B) Heatmap of metamodule correlations with clinical metadata. (C) Integration of cell-type specific pseudobulk DE data for each metamodule using kME threshold ≥0.75 with non-asthma set as reference. Percent of genes from each cell-type meeting threshold for adjusted p-value significance and represented in the top 20 enriched genes in each metamodule denoted. FEV1 – forced expiratory volume in one second, Macs – macrophages, Eos – eosinophils, BSA – body surface area, Prevent 10/30 – cardiovascular risk score in 10/30 years as calculated by PREVENT algorithm, Hb A1c - % glycosylated hemoglobin, VAT – visceral adipose tissue, BRI – body roundness index, W/H – waist-to-hip, BMI – body mass index, FMI – fat mass index, TG/HDL – triglycerides-to-high density lipoprotein, CRP – high sensitivity c-reactive protein, HOMA-IR-hemostatic measure of insulin resistance, cOCS – courses of oral corticosteroid, Dx – diagnosis, FeNO – fraction exhaled nitric oxide, ICS – inhaled corticosteroid dose. * P<0.05; ** P<0.01; *** P<0.001; **** p<0.0001

E16 was the largest module in the asthma-associated metamodule E (Figure 5A). It encompassed a broad activation signature, collectively suggesting integration of bioenergetic activity, protein synthesis, structural remodeling, and effector immune function (Supplemental Table 1). Clinically, module E16 was positively associated with asthma disease activity and severity, as seen by positive correlation with exacerbation frequency, ICS dose and disease duration and inverse correlation with lung function and sputum macrophages. Module E16 revealed a strong upregulation in CD4⁺ memory T cells and more modest increases in CD8⁺ memory T cells, mature NK cells and Tregs based on asthma diagnosis (Figure 5C). This pattern supports a model in which mitochondrial and transcriptional activation programs are amplified in asthma primarily within CD4⁺ TEM cells, with secondary engagement of additional adaptive and cytotoxic lymphocyte subsets.

Modules E13 and E14 had strong interactions with other modules, including E16 (Figure 5A). E13 and E14 were enriched for pathways reflecting mitochondrial activity, protein synthesis and turnover, organelle dynamics, and cellular stress adaptation. Together, these modules define a metabolically activated cellular state engaged in protein synthesis consistent with heightened bioenergetic demand and stress signaling. Clinically, modules E13 and E14 were positively associated with IgE levels and inversely correlated with FEV1. E13 additionally associated with increased blood neutrophil counts, whereas E14 showed broader links to asthma disease activity and severity, including positive associations with FeNO, exacerbation frequency, ICS dose, and older age at diagnosis, alongside reduced sputum macrophage proportions. No single cell population uniquely accounted for the module E13 gene expression with modest contribution from CD4⁺ TEM in module E14 (Figure 5C).

Modules E17-E19 defined a coordinated axis of interferon signaling, T-cell activation, and T2 immune programming associated with asthma and disease severity. Clinically, modules E17-E19 were positively associated with exacerbation frequency, ICS dose, bronchodilator reversibility, and/or changes in FEV1, consistent with impaired lung function. Collectively, modules E17-E19 integrated innate immune activation, chemokine-mediated recruitment, adaptive T-cell stimulation, and stress-adaptive survival pathways. E17 was enriched for interferon-responsive and innate immune genes alongside immune activation markers in CD4⁺ TEM and Treg. E18 reflected a T2 immune program centered on GATA3, a master regulator of Th2 differentiation, and a strong upregulation of CD4⁺ TEM cells. E19 further emphasized adaptive immune activation and cellular persistence and included genes involved in T-cell stimulation and co-signaling, immune regulation and migration, and protein synthesis. Modules C7 through D12 (encompassing all metamodules C and D) were correlated with non-asthma status and enriched for pathways related to transcriptional regulation, RNA processing, protein trafficking, genome stability, and immune modulation. Together, these modules define a coordinated regulatory network supporting cellular adaptation and homeostasis that is absent in AT immune cells in asthma.

### Asthma-specific T/NK Cell iterative WGCNA identify modules with a T2 inflammatory signature

Since asthma is a heterogeneous inflammatory disease with many inflammatory phenotypes, we repeated the iterativeWGCNA using only the asthma cohort (n=16) T/NK cells (Figure 6A-B). Four dominant clinically defined clusters of modules emerged from the co-expression analysis. Modules S71-73 (Light Blue) plus T75 and Q64 identified a cluster of modules that were positively correlated with classical biomarkers of T2 inflammation (blood eosinophil count, % sputum eosinophils, and serum periostin), but not FeNO. Genes upregulated in S71-73 are involved in metabolic stress signaling, extracellular matrix remodeling, chromatin and transcriptional regulators, long non-coding RNAs, stress response and protein synthesis genes, and regulators of proliferation and signaling (Supplemental Table 2). Collectively, modules S71-73 suggest a coordinated stress and tissue remodeling response linked to airway eosinophilia which operates independently of airway nitric oxide handling in obesity.

**Figure 6.**
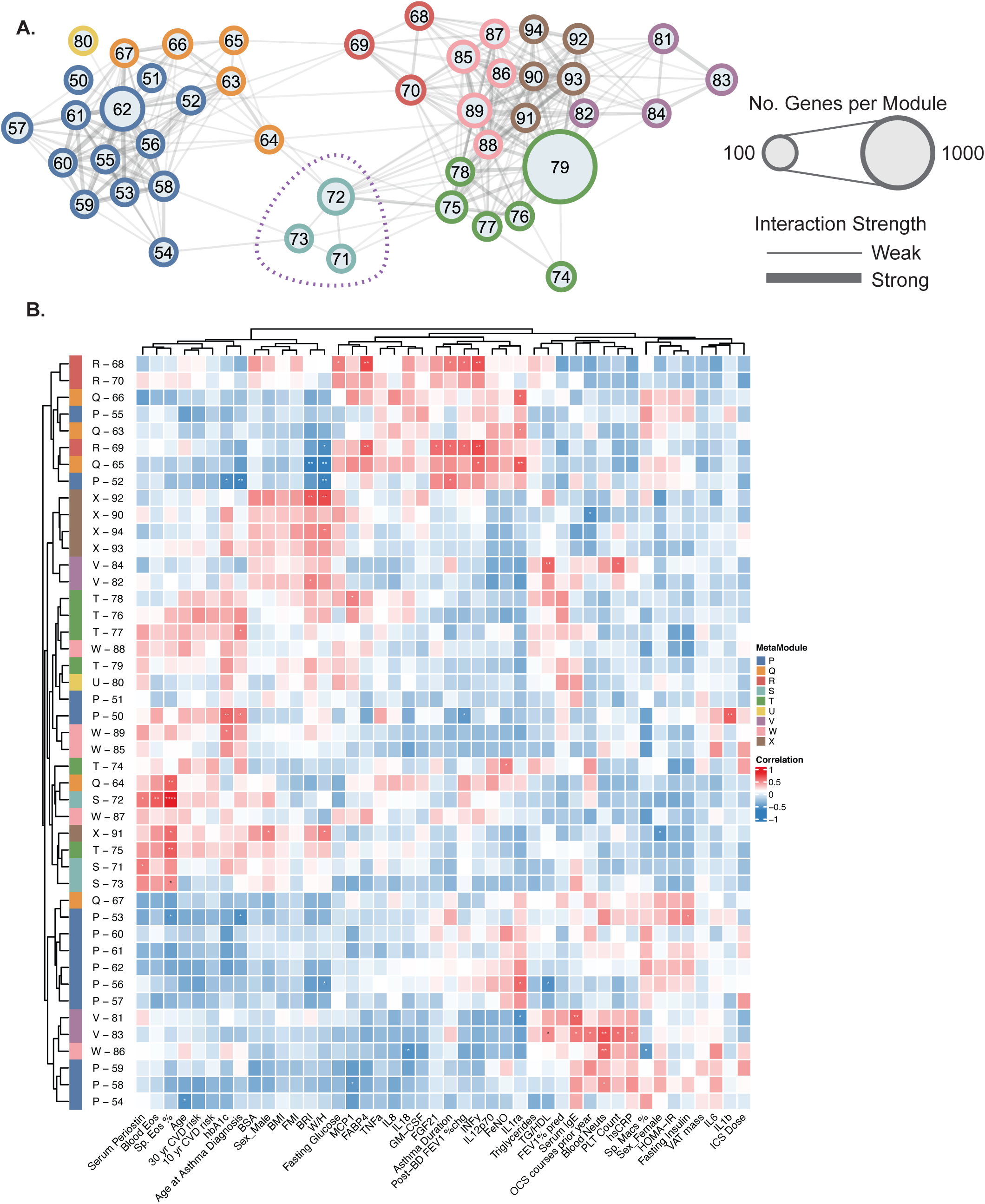
– Asthma-restricted iterativeWGCNA of T/NK fraction identifies distinct metamodules associated with T2 inflammation, systemic metabolism, and central adiposity. A) iterativeWGCNA module and metamodule network. B) Heatmap of metamodule correlations with clinical metadata. cOCS – courses of oral corticosteroid, ICS – inhaled corticosteroid dose, FeNO – fraction exhaled nitric oxide, PLT – platelet count, TG/HDL – triglycerides-to-high density lipoprotein, HOMA-IR-hemostatic measure of insulin resistance, FM – fat mass, VAT – visceral adipose tissue, BSA – body surface area, BRI – body roundness index, W/H – waist-to-hip, CVD – cardiovascular disease, Sp – sputum, FEV1 – forced expiratory volume in one second, post-BD – post-bronchodilator. * P<0.05; ** P<0.01; *** P<0.001; **** p<0.0001

Modules V81-84 (Purple) and X90-94 (Brown) positively correlated with systemic inflammation (platelet count, blood neutrophils, and hsCRP) and/or central adiposity (waist-to-hip ratio and body roundness). Serum total IgE and exacerbation frequency also correlated with modules V81 and 83 which were further correlated with systemic inflammation. Modules R68-70 (Red) correlated with IFNγ and FABP4 while modules Q63-Q65 (Orange) correlated with IL-1RA levels. Metamodules P (Blue) and T (Green) had few associations with any of the clinical parameters available in our dataset. Interestingly, serum IL-6 levels did not show a strong correlation with any module in the T/NK set. This supports that there are multifactorial processes at play in obesity-associated asthma that contribute to heightened AT inflammation and dysfunction, which are anticipated to extend beyond classical clinical and biomarker indicators of asthma and metabolic dysfunction.

### Myeloid iterativeWGCNA shows that asthma diagnosis is the driving factor for module formation

A similar iterativeWGCNA analysis was conducted on the myeloid cell fraction with asthma status as the main determining factor of co-expression (Figure 7A-B) with genes in specific cell populations for each metamodule shown in Supplemental Figure 3. These data reiterated that asthma status in obesity influences AT immune gene correlations more than any other clinical parameter (Figure 7B). Five major metamodules (AA–EE) were identified, each capturing distinct biological programs and clinical relationships. Metamodule AA showed limited cell type enrichment and was associated with type 2 inflammatory features. In contrast, metamodule BB, enriched in monocyte and macrophage populations, was downregulated in asthma and instead associated with metabolic parameters (fasting glucose, VAT mass and IL-8), suggesting a macrophage-driven immunoregulatory program suppressed in asthma and associated with the loss of airway macrophages. Module B13, which is not specific to any cell type, stands out as the sole module across all iterativeWGCNA analyses highly associated with serum IL-6. Metamodules CC, DD, and EE were all upregulated in asthma but reflected distinct pathological axes. Metamodule CC captured a translational and oxidative phosphorylation program, enriched in LAMs and PVMs, associated with disease severity and reduced lung function (Supplemental Table 3). Metamodule DD showed the strongest linkage to classical monocytes and was tightly associated with asthma diagnosis, severity, exacerbations, and type 2 biomarkers, with modules DD27–30 defining a coordinated allergic inflammatory network. Metamodule EE represented a lipid metabolism–driven macrophage program linked to both adiposity and asthma severity, integrating metabolic dysfunction with immune activation. Body composition, other inflammatory biomarkers, and cardiometabolic clinical parameters showed minimal association with module and metamodule formation.

**Figure 7.**
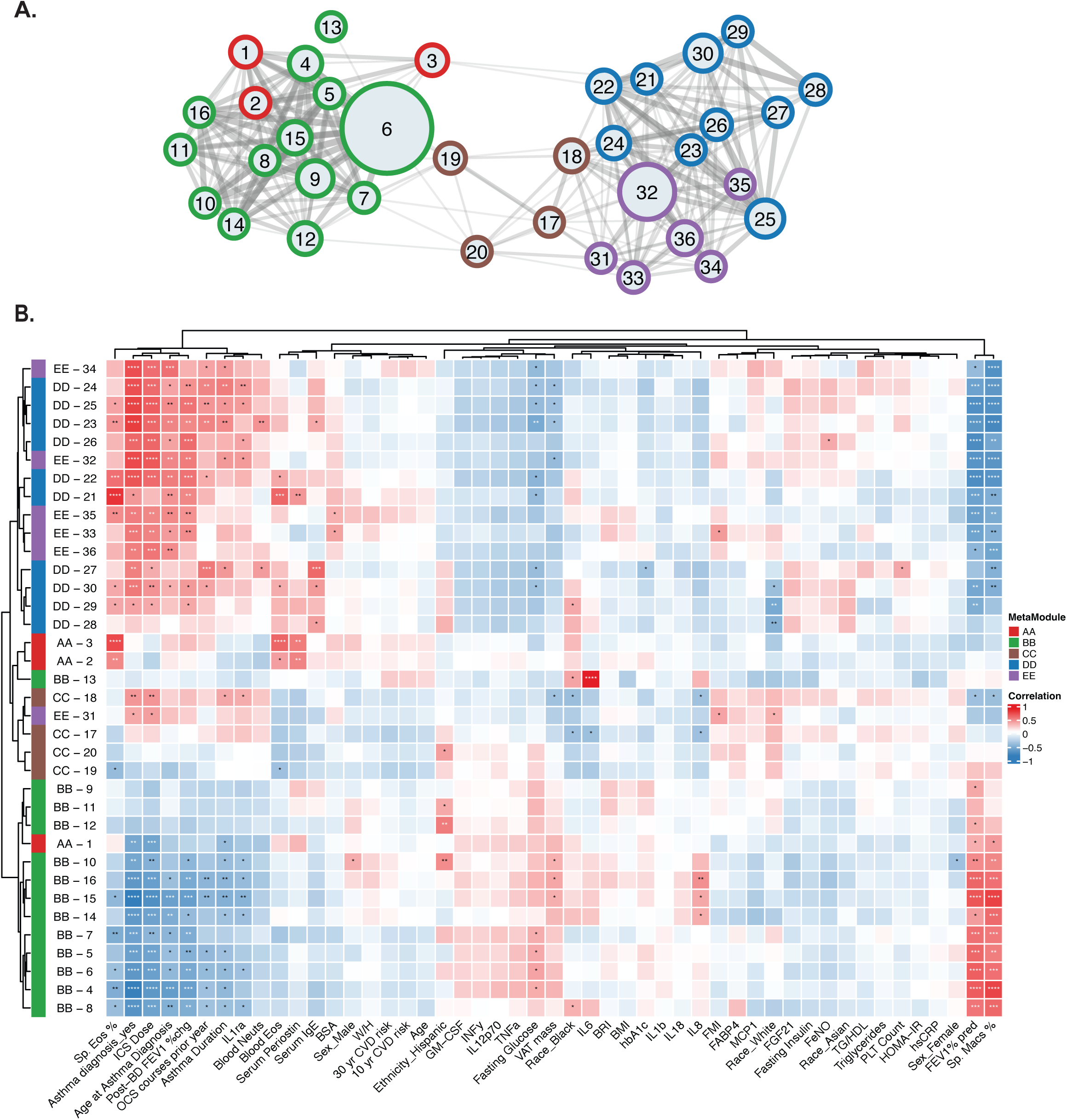
– IterativeWGCNA of myeloid fraction identifies asthma as the clinical variable driving module formation. A) iterativeWGCNA module and metamodule network. B) Heatmap of metamodule correlations with clinical metadata. cOCS – courses of oral corticosteroid, ICS – inhaled corticosteroid, FeNO – fraction exhaled nitric oxide, PLT – platelet, TG/HDL – triglyceride-to-high density lipoprotein, HOMA-IR – hemostatic model of insulin resistance, FM – fat mass, VAT – visceral adipose tissue, BSA – body surface area, BRI – body roundness index, W/H – waist-to-hip, CVD – cardiovascular risk, FEV1 – forced expiratory volume in one second, pred – predicted, post-BD – post-bronchodilator response, sp – sputum. * P<0.05; ** P<0.01; *** P<0.001; **** p<0.0001

## Discussion

Obesity is both a risk factor and a modifier of risk for many chronic inflammatory conditions. Asthma, a disease of chronic airway inflammation, is a classic example of a clinical disease state with clear association to and modification by excess adipose mass. In the context of asthma, excess fat mass is thought to influence the lung through direct mass effects on lung mechanics, systemic inflammatory mediators such as IL-6 and adipokines which may augment airway inflammation, and the convergence of lifestyle factors such as diet macro- and micro-nutrient composition, physical activity, and microbiome alterations on the airway. Conversely, asthma predisposes to obesity development and cardiometabolic risk with lifestyle factors and corticosteroid exposures implicated (57–60). Utilizing single-cell sequencing of AT immune cells from individuals with obesity and/or without clinical asthma and similar levels of excess fat mass, metabolic health, systemic proinflammatory mediators, age, race and sex, we identified asthma status alone as a key mediator of a novel asthma-associated AT inflammatory signal. AT from individuals with obesity-associated asthma is marked by increased LAMs and PVMs, decreased naïve T cells, increased metabolic activity in naïve and effector cells and macrophages, and a dysfunctional activation signature. Asthma diagnosis is the key clinical factor observed in the iterativeWGCNA which correlated with the differences in the T/NK and myeloid cell immune signatures. Our data represents the first evidence of a chronic respiratory disease associated with enhanced inflammation in the AT and supports that asthma, in the context of obesity, is a chronic inflammatory disorder associated with an inflammatory state that extends beyond the airway to involve distant tissues including the adipose.

While this cross-sectional study cannot confirm the directionality of the relationship between AT immune changes and asthma status, the presence of classical Th2 and Th17 T cell signatures, the correlation between classical biomarkers of the T2-high asthma phenotype, the dominant asthma phenotype in the absence of obesity, and a restricted set of modules in the asthma-only T/NK cell and myeloid iterativeWGCNA, support the hypothesis that classical airway inflammatory signatures extend to the AT and influence immune cell profiles in AT. Inhaled corticosteroid dose and oral corticosteroid use in the prior one year, the available surrogates of corticosteroid exposure in the asthma cohort, showed variable correlation with modules in the T/NK and myeloid iterativeWCGNA, and minimal correlation in the asthma-only iterativeWGCNA implying the pathobiology of asthma, and not the drugs we use to treat asthma, correlates with many of the unique AT signatures observed in the context of asthma. Additionally, the separation of T2-high modules from those correlated with central adiposity and biomarkers of systemic inflammation support the complexity of clinical features which augment inflammatory state in AT in the setting of obesity and comorbid asthma. Profiling of AT from lean individuals with asthma and individuals on a variety of asthma treatment strategies would confirm the impact of T2 airway inflammation on AT immune dysregulation.

Interestingly, the enhanced AT inflammation we observed in the context of asthma was not mirrored in the assessed systemic inflammatory biomarkers, including clinically available hsCRP and IL-6. Biomarker assessment was particularly targeted towards macrophage and T1 inflammatory pathways which may be of greatest relevance to AT inflammation in the obese state. IL-1RA is a notable exception but may not reflect an inflammatory AT signature. IL-1RA protein levels are elevated in pediatric patients with acute asthma exacerbation, polymorphisms in the *IL1RA* are associated with risk for asthma development, and IL-1RA serves a protective role at the airway epithelium during viral infection with RSV (61–63). Similarly, elevations in serum MCP-1 are reported in asthma cohorts and increase with exacerbations(64).The limited increased systemic inflammatory biomarkers in asthma further suggests a distinct airway-AT inflammatory axis that may not otherwise be detected through classical systemic assessments. Paracrine mechanisms, which would be particularly relevant from adipose depots surrounding the large airway, cannot be excluded as an unobserved mechanism in our study (65).

Impaired response to airway restricted (inhaled corticosteroids) and systemic asthma therapies (systemic corticosteroids, targeted asthma biologic therapies) are well established in obesity-associated asthma (7–9). Our findings of enhanced AT inflammation and immune cell dysfunction support that AT may be a mechanism of resistance, serving as a depot of immune cells to propagate therapeutic resistance. Studies which evaluate the effect of systemic therapy on AT immune dysregulation may unlock critical mechanisms underlying this clinical problem. Additionally, asthma specific AT immune dysregulation may have implications for disease persistence and the newer clinical concept of asthma remission. Resolution of local airway inflammation may be insufficient to cause remission of asthma in the context of obesity. Studies of patients with asthma undergoing bariatric surgery support that large reductions in AT mass may lead to asthma remission (66).

## Conclusion

Adipose tissue in the context of obesity-associated asthma is marked by increased myeloid and T effector cell abundance and increased activation state with features of immune cell dysfunction. Asthma differentially affects immune cell frequency and metabolism in AT. Asthma in the context of obesity represents a systemic inflammatory process with associated AT tissue inflammation that is not captured by conventional systemic inflammatory biomarkers. This has critically important potential therapeutic implications for patients with asthma that need to be rigorously studied.

## Methods

### Patient cohorts and clinical metadata

Patients of similar age, sex, race, ethnicity, BMI, A1C levels, and PREVENT score risk for 10-and 30-year cardiovascular risk were selected from two existing cohorts (N=16 each), GLP-1R Agonists in the Treatment of Adult, Symptomatic, Obese Asthma (GATA-3) and Cardiovascular Effects of GLP-1 Receptor Activation (CEGRA) studies. All patients had BMI >27 kg/m2 and an A1C level that was below the threshold for type 2 diabetes. In the GATA-3 cohort, asthma was defined as moderate-to-severe persistent based on a requirement for medium-dose inhaled corticosteroid or more therapy. Asthma status was confirmed by evidence of bronchodilator response or methacholine-induced airways hyperresponsiveness. Oral glucocorticoid use in the 56 days prior to sampling and asthma biologic use in the prior 5 months were exclusions. Patients in the CEGRA cohort were excluded if asthma diagnosis was noted on chart review. Our study examined adipose tissue from males and females and biologic sex was included in our matching and analysis strategies.

### Subcutaneous adipose tissue sampling

Abdominal subcutaneous AT biopsies were performed approximately 3 cm to the right of the umbilicus as previously described (67). Briefly, skin and AT were anesthetized with 1% lidocaine, some with 1:100,000 epinephrine, and approximately 5 grams of AT were collected using a 2.1 mm blunt, side-ported liposuction catheter into saline. Visible clots were removed, the AT was strained using a 70um filter, and tissue was separated into a single cell suspension using gentleMACS Dissociator and collagenase. Single cell suspensions were then crypreserved for downstream use.

### Inflammatory Biomarker Analysis

Serum samples were analyzed for IL-6 on the Roche cobas® e 801 module and hsCRP on the Roche cobas® 4000 analyzer through the VUMC Laboratory for Translational and Clinical Research. Citrate plasma samples were analyzed on Meso Scale Discovery© human platforms for FABP4, leptin, FGF-21, GM-CSF, IFN-γ, IL-1β, IL-1RA, IL-8, IL-12p70, IL-18, MCP-1, and TNF-α.

### Single cell RNA sequencing

scRNA-seq was conducted on the stromovascular fractions of subcutaneous AT. Five thousand cells were targeted for sequencing from each sample using the single-cell capturing and library construction Chromium Single-Cell 3’ Reagent Kits (v3.1 chemistry, PN-1000130) from 10x Genomics according to the manufacturer’s instructions. scRNA-seq libraries were constructed using 10X Genomics Library Construction Kits (PN-1000196) and Dual Index Kit TT Set A (PN-1000215). The libraries were paired-end sequenced on the Illumina NovaSeq X.

### Bioinformatic and statistical analysis

#### Single cell RNA-Seq analysis

Raw sequencing reads were aligned to the 10X GRCh38 human reference genome (refdata-gex-GRCh38-2020-A) using Cell Ranger v9.0.0 (68). Ambient RNA contamination was detected and removed with CellBender v0.3.2 (69, 70). Per-sample CellBender-filtered HDF5 matrices were loaded into R v4.3.2 using Seurat v5.1.0 (70), retaining features detected in at least 3 cells and cells with at least 200 detected features. Per-cell quality metrics were calculated, including the percentage of counts mapping to mitochondrial (^MT-), small ribosomal subunit (^RPS), and large ribosomal subunit (^RPL) genes. Cells were filtered in two passes: first by a sample-specific UMI threshold (retaining cells within mean ± 2 SD of each sample’s total UMI distribution), then by global hard cutoffs (nFeature_RNA: 250–5,000; nCount_RNA: 1,000–35,000; percent mitochondrial: < 20%). Filtered samples were merged and processed with log normalization, selection of 2,000 highly variable features (variance-stabilizing transformation), scaling, and PCA (50 components). A shared nearest-neighbor graph was constructed on the top 30 PCs and cells were clustered using the Louvain algorithm (resolution 0.4). Batch effects attributable to processing library and experimental batch were used as covariates during integration with Harmony v1.2.3(71). Doublets were identified using the cluster-based mode of scDblFinder v1.16.0 (72), applied per sample using Harmony-integrated cluster assignments at resolution 0.6; roughly 11% of cells were classified as doublets and excluded. Singlets were re-normalized and re-processed (2,000 HVGs, 20 PCs, Harmony correction on library and batch, UMAP with 20 PCs, Louvain resolution 0.4). Coarse cell type labels were assigned to clusters by canonical marker gene expression curated from Bailin et al.(28). The myeloid and T/NK compartments were each independently subsetted and subjected to iterative reclustering with Harmony integration (20 PCs; Louvain resolution 1.0 for myeloid cells; resolution optimized per compartment for T/NK cells). Clusters representing cross-lineage doublets were identified by marker expression and removed. To improve resolution of important immune cell subgroups, we performed reclustering on the T/NK cell and myeloid cell fractions. The T/NK cell and myeloid cell clusters were utilized for all subsequent downstream analyses. Final myeloid annotations comprised: classical monocytes, non-classical monocytes, intermediate monocytes, lipid-associated macrophages (LAM), perivascular macrophages, cDC1, cDC2B, and migratory DCs. Final T/NK annotations comprised: CD4 memory T cells, CD4 naive T cells, CD8 memory T cells, CD8 naive T cells, NK cells, and MAST cells.

### Compositional Analysis of Cell Type Abundances

Changes in cell type composition between asthma-diagnosed and healthy donors were assessed using scCODA v0.1.9 (29). Raw cell type counts aggregated per donor were used as input, separately for the myeloid and T/NK compartments. A single covariate (condition: asthma_diagnosed vs. no asthma) was included in the model formula. The reference cell type for each compartment was selected as the cell type with the lowest median log-ratio standard deviation across samples, an approach that identifies the most compositionally stable population (cDC2B for myeloid; CD8 memory T cells for T/NK). Posterior inference was performed using the No-U-Turn Sampler (NUTS) implemented via numpyro v0.18.0 and JAX v0.6.0, with 11,000 sampling iterations and a random seed of 1234. Cell type abundance changes were considered credible at a false discovery rate of 0.05, defined as the posterior probability that the highest-density interval of the effect estimate excludes zero. Sensitivity analyses were additionally performed at FDR = 0.1

### Pseudobulk Differential Expression Analysis

Pseudobulk count matrices were generated separately for the myeloid and T/NK compartments, both at the whole-compartment level and stratified by annotated cell type, using Seurat’s AggregateExpression. Genes were retained if the sum of raw counts across all pseudobulk samples exceeded zero and further filtered to genes with DESeq2-normalized counts ≥ 5 in at least 3 samples. Principal component analysis of VST-transformed data was performed to assess potential confounders between principal components and technical/clinical covariates (age, BMI, HbA1c, sex, race, ethnicity, library, and batch). Differential expression between asthma-diagnosed and healthy donors was modeled using DESeq2 v1.42.1(73) with the design formula ∼ sex + condition. All reported log₂ fold changes and adjusted p-values (Benjamini–Hochberg) are from the DESeq2 Wald test.

### Gene Set Enrichment Analysis

Pre-ranked GSEA was performed using the DESeq2 Wald test statistic as the ranking metric. Gene sets were retrieved from MSigDB (74) via msigdbr v25.1.1 (75) for three collections: Hallmark (H), curated gene sets (C2, C7, including KEGG), and ontology gene sets (C5). Genes with missing Wald statistics were excluded prior to ranking. GSEA was executed with clusterProfiler v4.10.1 (76) at an adjusted p-value cutoff of 0.05 and run independently for each annotated cell type within both the myeloid and T/NK compartments.

### Co-expression Network Analysis

We applied iterativeWGCNA (77) with an eigengene connectivity (kME) value of 0.75, minimum module size of 40 genes for biological relevance, and cell type-specific soft-thresholding power (power of 6 for T/NK cells, power of 8 for myeloid cells, power of 8 for T/NK cells/asthma-only samples, power of 10 for myeloid cells/asthma-only samples). Existing clinical metadata available for each cohort was incorporated where available. Missing data was imputed where applicable for the no asthma cohort. Specifically, metabolic variables including glucose, insulin, IL-6, hsCRP, total fat mass percentage, trunk fat mass percentage, and visceral adipose tissue (VAT) mass were imputed using the median values calculated across the combined study population, to preserve overall distributional properties. For derived anthropometric and cardiometabolic indices (fat mass index, body surface area, body roundness index, waist-to-hip ratio, triglycerides, triglyceride-to-HDL ratio, blood neutrophil count, and platelet count), median values from the asthma cohort were applied to the non-asthma cohort. These variables are primarily driven by adiposity and metabolic status rather than asthma per se, and both cohorts were matched for obesity; therefore, asthma cohort medians were used to preserve biologically relevant distributions while maintaining internal consistency among interdependent features. Serum periostin, for which population-level reference data are limited, was similarly imputed using the asthma cohort median to enable inclusion in integrative analyses while avoiding extrapolation beyond observed ranges. For variables not directly measured in the non-asthma cohort, imputation was guided by population-based reference data and literature-derived distributions. Blood eosinophil counts were imputed based on distributions reported in NHANES study (78). Total IgE levels and percent predicted FEV1 were assigned using values derived from published cohort data, including obese populations without asthma(79), selected to reflect physiologically plausible, non-asthmatic ranges. Sputum eosinophil and macrophage proportions were estimated using reference values from sputum inflammatory cell study(80), representing typical airway cellular distributions in non-asthmatic populations.

Networks were constructed using signed correlation preserving positive/negative correlations and maintaining biological directionality. Pseudobulk gene expression data derived from the scRNA-Seq data were variance-stabilized using DESeq2 and filtered to retain genes with expression above the median library size divided by 1 million (approximately 18 counts per million reads based on median library size of ∼17.8 million reads) in at least 5 samples per condition, resulting in 10,734 genes. iterativeWGCNA was used to construct a meta-network and estimate topological overlap among the modules based on correlations between detected module eigengenes. The Spinglass community detection algorithm, as implemented in the iGraph library (81), was used to group modules with a large degree of topological overlap into meta-modules (parameters: correlation threshold of 0.5, gamma of 1.9). Hierarchical clustering (correlation-based) of modules within meta-modules was performed on their eigengenes. The module meta-network was visualized using Cytoscape (82).

Enriched pathways were identified using the package clusterProfiler in R. Enriched gene functions were identified using gene ontology (GO) annotations obtained from the appropriate species-specific Bioconductor annotation database (org.Hs.eg.db) and KEGG pathway (83) analysis using clusterProfiler. The complete set of expressed genes was used as the reference set in all enrichment analyses with a p-value cutoff of 0.05 and q-value cutoff of 0.2. Minimum gene set size was set to 5 genes for pathway analysis.

Transcription factors were identified as genes annotated with the GO term ‘transcription regulator activity’ (GO:0140110) obtained from AmiGO. Genes involved in signaling pathways were identified as those annotated with ‘signal transduction’ (GO:0007165). Long non-coding RNAs were identified from GENCODE annotations using GTF files (84). Enrichment of each gene set within individual modules was evaluated using Fisher’s exact test implemented in the iterativeWGCNA pipeline.

### Statistical Analysis

Descriptive metadata reported as mean ± standard deviation where normally distributed. Wilcoxon test compared biomarker values between disease state. Data was analyzed using GraphPad Prism 10 for Windows version 10.6.1, ©GraphPad Software, LLC.

## Study Approval

This study was approved by the local institutional review board and conducted according to Declaration of Helsinki principles. Written informed consent was obtained from all study participants before any study participation and all study procedures.

## Data availability

For the single cell RNA-Seq data described in this publication, the raw data, processed files, and single cell objects have been deposited in NCBI’s Gene Expression Omnibus (GEO) and are available under accession number GSE327589. Additional metadata information is available directly from the corresponding author.

## Author contributions

DCN, AT, MHB, SB, JRK, KNC, MM designed the research study, DN, AT, BIS, LH, SH, KNC and MM conducted experiments and acquired data. J-PC, SS, AT, KNC, DN, MM analyzed the data. KN, JML, NJB, MM, KNC provided reagents, KNC, DN, and AT wrote the first draft of the manuscript. All authors reviewed and revised the manuscript.

Co-first authorship order was determined based on duration of contribution to the lifecycle of the research project.

## References

1. Ward ZJ, et al. Projected U.S. State-Level Prevalence of Adult Obesity and Severe Obesity. N Engl J Med. 2019;381(25):2440–2450.

2. Bloodworth MH, et al. Impact of metabolic and weight components on incident asthma using a real-world cohort. Ann Allergy Asthma Immunol. 2024;S1081-1206(24)01509–6.

3. Beuther DA, Sutherland ER. Overweight, obesity, and incident asthma: a meta-analysis of prospective epidemiologic studies. Am J Respir Crit Care Med. 2007;175(7):661–666.

4. Chen YC, et al. Gender difference of childhood overweight and obesity in predicting the risk of incident asthma: a systematic review and meta-analysis. Obes Rev. 2013;14(3):222–231.

5. Egan KB, Ettinger AS, Bracken MB. Childhood body mass index and subsequent physician-diagnosed asthma: a systematic review and meta-analysis of prospective cohort studies. BMC Pediatr. 2013;13:121.

6. Parasuaraman G, et al. The association between body mass index, abdominal fatness, and weight change and the risk of adult asthma: a systematic review and meta-analysis of cohort studies. Sci Rep. 2023;13(1):7745.

7. Gonem S, et al. Effects of Obesity on Response to Asthma Biologic Treatment: Longitudinal Data From the United Kingdom Severe Asthma Registry. The Journal of Allergy and Clinical Immunology: In Practice. 2025;13(11):3002–3010.

8. Peters-Golden M, et al. Influence of body mass index on the response to asthma controller agents. Eur Respir J. 2006;27(3):495–503.

9. Boulet L-P, Franssen E. Influence of obesity on response to fluticasone with or without salmeterol in moderate asthma. Respir Med. 2007;101(11):2240–2247.

10. Peters MC, et al. Plasma interleukin-6 concentrations, metabolic dysfunction, and asthma severity: a cross-sectional analysis of two cohorts. Lancet Respir Med. 2016;4(7):574–584.

11. Peters MC, et al. The Impact of Insulin Resistance on Loss of Lung Function and Response to Treatment in Asthma. Am J Respir Crit Care Med. 2022;206(9):1096–1106.

12. Vijay J, et al. Single-cell analysis of human adipose tissue identifies depot- and disease-specific cell types. Nat Metab. 2020;2(1):97–109.

13. Englebert K, et al. The CD27/CD70 pathway negatively regulates visceral adipose tissue-resident Th2 cells and controls metabolic homeostasis. Cell Rep. 2024;43(3):113824.

14. Yesian AR, et al. Preadipocyte IL-13/IL-13Rα1 signaling regulates beige adipogenesis through modulation of PPARγ activity. J Clin Invest. 2025;135(11):e169152.

15. Brestoff JR, et al. Group 2 innate lymphoid cells promote beiging of white adipose tissue and limit obesity. Nature. 2015;519(7542):242–246.

16. Wu D, et al. Eosinophils Sustain Adipose Alternatively Activated Macrophages Associated with Glucose Homeostasis. Science. 2011;332(6026):243–247.

17. Jacks RD, Lumeng CN. Macrophage and T cell networks in adipose tissue. Nat Rev Endocrinol. 2024;20(1):50–61.

18. Chavakis T, Alexaki VI, Ferrante AW. Macrophage function in adipose tissue homeostasis and metabolic inflammation. Nat Immunol. 2023;24(5):757–766.

19. Kim HY, et al. Interleukin-17–producing innate lymphoid cells and the NLRP3 inflammasome facilitate obesity-associated airway hyperreactivity. Nat Med. 2014;20(1):54–61.

20. Toki S, et al. Glucagon-like peptide-1 receptor agonist inhibits aeroallergen-induced activation of ILC2 and neutrophilic airway inflammation in obese mice. Allergy. 2021;76(11):3433–3445.

21. Bapat SP, et al. Obesity alters pathology and treatment response in inflammatory disease. Nature. 2022;604(7905):337–342.

22. Newcomb DC, et al. Estrogen and progesterone decrease let-7f microRNA expression and increase IL-23/IL-23 receptor signaling and IL-17A production in patients with severe asthma. J Allergy Clin Immunol. 2015;136(4):1025–1034.e11.

23. Chowdhury NU, et al. Androgen signaling restricts glutaminolysis to drive sex-specific Th17 metabolism in allergic airway inflammation. J Clin Invest. 2024;134(23):e177242.

24. McLaughlin T, et al. T-cell profile in adipose tissue is associated with insulin resistance and systemic inflammation in humans. Arterioscler Thromb Vasc Biol. 2014;34(12):2637–2643.

25. Li X, et al. Investigation of the relationship between IL-6 and type 2 biomarkers in patients with severe asthma. Journal of Allergy and Clinical Immunology. 2020;145(1):430–433.

26. Ohsugi M, et al. Real-world use of glucagon-like peptide-1 receptor agonists in Japanese patients with type 2 diabetes: A retrospective database study (DEFINE-G). Diabetes Res Clin Pract. 2023;203:110841.

27. Global Initiative for Asthma. Global Strategy for Asthma Management and Prevention [Internet]. 2025. https://www.ginasthma.org.

28. Bailin SS, et al. Changes in subcutaneous white adipose tissue cellular composition and molecular programs underlie glucose intolerance in persons with HIV. Front Immunol. 2023;14:1152003.

29. Büttner M, et al. scCODA is a Bayesian model for compositional single-cell data analysis. Nat Commun. 2021;12(1):6876.

30. Lan F, et al. GZMK-expressing CD8+ T cells promote recurrent airway inflammatory diseases. Nature. 2025;638(8050):490–498.

31. Zhang H, et al. Dysfunction of S100A4+ effector memory CD8+ T cells aggravates asthma. European Journal of Immunology. 2022;52(6):978–993.

32. Pan Y, et al. Survival of tissue-resident memory T cells requires exogenous lipid uptake and metabolism. Nature. 2017;543(7644):252–256.

33. Ferry A, et al. The XCL1-XCR1 axis supports intestinal tissue residency and antitumor immunity. J Exp Med. 2025;222(2):e20240776.

34. Ward-Kavanagh LK, et al. The TNF Receptor Superfamily in Co-stimulating and Co-inhibitory Responses. Immunity. 2016;44(5):1005–1019.

35. Ramonell RP, et al. CD8+ TEMRAs in severe asthma associate with asthma symptom duration and escape proliferation arrest. JCI Insight. 2025;10(8):e185061.

36. van der Ploeg EK, et al. Type-2 CD8+ T-cell formation relies on interleukin-33 and is linked to asthma exacerbations. Nat Commun. 2023;14(1):5137.

37. Bruchard M, et al. The receptor NLRP3 is a transcriptional regulator of TH2 differentiation. Nat Immunol. 2015;16(8):859–870.

38. Kumar R, Theiss AL, Venuprasad K. RORγt protein modifications and IL-17-mediated inflammation. Trends in Immunology. 2021;42(11):1037–1050.

39. Lin R, et al. GPR65 promotes intestinal mucosal Th1 and Th17 cell differentiation and gut inflammation through downregulating NUAK2. Clin Transl Med. 2022;12(3):e771.

40. Katsouda A, et al. MPST sulfurtransferase maintains mitochondrial protein import and cellular bioenergetics to attenuate obesity. Journal of Experimental Medicine. 2022;219(7):e20211894.

41. Zou Q, Qi H. Deletion of ribosomal paralogs Rpl39 and Rpl39l compromises cell proliferation via protein synthesis and mitochondrial activity. Int J Biochem Cell Biol. 2021;139:106070.

42. Yánez DC, et al. IFITM proteins drive type 2 T helper cell differentiation and exacerbate allergic airway inflammation. Eur J Immunol. 2019;49(1):66–78.

43. Costa-Pereira S, et al. Regulatory T cells suppress GM-CSF-producing T helper cells via IL-2 modulation to restrain immunopathology. Cell Reports. 2025;44(5):115642.

44. Pan Y, et al. Survival of tissue-resident memory T cells requires exogenous lipid uptake and metabolism. Nature. 2017;543(7644):252–256.

45. Singer M, et al. A Distinct Gene Module for Dysfunction Uncoupled from Activation in Tumor-Infiltrating T Cells. Cell. 2016;166(6):1500–1511.e9.

46. Karin M, et al. Metallothionein mRNA induction in HeLa cells in response to zinc or dexamethasone is a primary induction response. Nature. 1980;286(5770):295–297.

47. Kim CH, et al. Zinc-induced NF-kappaB inhibition can be modulated by changes in the intracellular metallothionein level. Toxicol Appl Pharmacol. 2003;190(2):189–196.

48. Lietzau S, et al. Differential regulation of 5-lipoxygenase and leukotriene-C4-synthase expression by IFNγ, IL-4 and IL-13 in human monocytes and macrophages from patients with atopic dermatitis. Inflamm Res. 2025;74(1):175.

49. Chavey C, et al. CXC ligand 5 is an adipose-tissue derived factor that links obesity to insulin resistance. Cell Metab. 2009;9(4):339–349.

50. Sokulsky LA, et al. A Critical Role for the CXCL3/CXCL5/CXCR2 Neutrophilic Chemotactic Axis in the Regulation of Type 2 Responses in a Model of Rhinoviral-Induced Asthma Exacerbation. J Immunol. 2020;205(9):2468–2478.

51. Gauthier M, et al. CCL5 is a potential bridge between type 1 and type 2 inflammation in asthma. J Allergy Clin Immunol. 2023;152(1):94–106.e12.

52. Al-Mossawi H, et al. Context-specific regulation of surface and soluble IL7R expression by an autoimmune risk allele. Nat Commun. 2019;10(1):4575.

53. Liu M, et al. Grb10 Promotes Lipolysis and Thermogenesis by Phosphorylation-Dependent Feedback Inhibition of mTORC1. Cell Metabolism. 2014;19(6):967–980.

54. Shang W, et al. Deletion of EP3 prostaglandin receptor in murine macrophages aggravates diet-induced obesity by suppressing SPARC. EMBO J. 2025;44(18):4962–4983.

55. Liu B, et al. Sparcl1 promotes nonalcoholic steatohepatitis progression in mice through upregulation of CCL2. J Clin Invest. 2021;131(20). 10.1172/JCI144801.

56. Hildreth AD, et al. Single-cell sequencing of human white adipose tissue identifies new cell states in health and obesity. Nat Immunol. 2021;22(5):639–653.

57. Rodrigues M, et al. Early childhood asthma and adiposity until adolescence: A prospective birth cohort study using body fat percentage. Pediatr Allergy Immunol. 2026;37(5):e70360.

58. Bloom CI, et al. Association of Dose of Inhaled Corticosteroids and Frequency of Adverse Events. Am J Respir Crit Care Med. 2025;211(1):54–63.

59. Valencia-Hernández CA, et al. Asthma and incident coronary heart disease: an observational and Mendelian randomisation study. Eur Respir J. 2023;62(5):2301788.

60. Stratakis N, et al. The Role of Childhood Asthma in Obesity Development: A Nationwide US Multicohort Study. Epidemiology. 2022;33(1):131–140.

61. Sheikh S, et al. Innate immune responses are increased in children with acute asthma exacerbation. Pediatr Allergy Immunol. 2024;35(6):e14173.

62. Pattaro C, et al. Association between interleukin-1 receptor antagonist gene and asthma-related traits in a German adult population. Allergy. 2006;61(2):239–244.

63. Morris SB, et al. Long-term alterations in lung epithelial cells after EL-RSV infection exacerbate allergic responses through IL-1β-induced pathways. Mucosal Immunol. 2024;17(5):1072–1088.

64. Chan C-K, et al. Sequential evaluation of serum monocyte chemotactic protein 1 among asymptomatic state and acute exacerbation and remission of asthma in children. J Asthma. 2009;46(3):225–228.

65. Elliot JG, et al. Fatty airways: implications for obstructive disease. Eur Respir J. 2019;54(6):1900857.

66. Boulet L-P, et al. Effect of bariatric surgery on airway response and lung function in obese subjects with asthma. Respiratory Medicine. 2012;106(5):651–660.

67. Wanjalla CN, et al. Adipose Tissue in Persons With HIV Is Enriched for CD4+ T Effector Memory and T Effector Memory RA+ Cells, Which Show Higher CD69 Expression and CD57, CX3CR1, GPR56 Co-expression With Increasing Glucose Intolerance. Front Immunol. 2019;10:408.

68. Zheng GXY, et al. Massively parallel digital transcriptional profiling of single cells. Nat Commun. 2017;8(1):14049.

69. CellBender removes technical artifacts from single-cell RNA sequencing data. Nat Methods. 2023;20(9):1285–1286.

70. Satija R, et al. Spatial reconstruction of single-cell gene expression data. Nat Biotechnol. 2015;33(5):495–502.

71. Korsunsky I, et al. Fast, sensitive and accurate integration of single-cell data with Harmony. Nat Methods. 2019;16(12):1289–1296.

72. Germain P-L, et al. Doublet identification in single-cell sequencing data using scDblFinder. F1000Res. 2022;10:979.

73. Love MI, Huber W, Anders S. Moderated estimation of fold change and dispersion for RNA-seq data with DESeq2. Genome Biol. 2014;15(12):550.

74. Liberzon A, et al. The Molecular Signatures Database Hallmark Gene Set Collection. Cell Systems. 2015;1(6):417–425.

75. Dolgalev I. msigdbr: MSigDB Gene Sets for Multiple Organisms in a Tidy Data Format. 2018;26.1.0.

76. Yu G, et al. clusterProfiler: an R Package for Comparing Biological Themes Among Gene Clusters. OMICS: A Journal of Integrative Biology. 2012;16(5):284–287.

77. Greenfest-Allen E, et al. iterativeWGCNA: iterative refinement to improve module detection from WGCNA co-expression networks [preprint]. 2017. 10.1101/234062.

78. Malinovschi A, et al. Exhaled nitric oxide levels and blood eosinophil counts independently associate with wheeze and asthma events in National Health and Nutrition Examination Survey subjects. Journal of Allergy and Clinical Immunology. 2013;132(4):821–827.e5.

79. Dixon AE, et al. Effects of obesity and bariatric surgery on airway hyperresponsiveness, asthma control, and inflammation. Journal of Allergy and Clinical Immunology. 2011;128(3):508–515.e2.

80. Belda J, et al. Induced sputum cell counts in healthy adults. Am J Respir Crit Care Med. 2000;161(2 Pt 1):475–478.

81. Csárdi G, Nepusz T. The igraph software package for complex network research. 2006. https://igraph.org.

82. Shannon P, et al. Cytoscape: A Software Environment for Integrated Models of Biomolecular Interaction Networks. Genome Res. 2003;13(11):2498–2504.

83. Kanehisa M, et al. KEGG for taxonomy-based analysis of pathways and genomes. Nucleic Acids Research. 2023;51(D1):D587–D592.

84. Mudge JM, et al. GENCODE 2025: reference gene annotation for human and mouse. Nucleic Acids Research. 2025;53(D1):D966–D975.

