## Supplemental Tables 1-2 for "Adipose tissue as a site of immune activation and dysfunction in individuals with obesity and asthma"

Supplemental Table 1:

**T/NK iterativeWGCNA Module Key Gene Sets**

| Module | Key genes | Clinical Correlation | Integrated interpretation |
|---|---|---|---|
| **B2** | Protein turnover and trafficking: *CMSS1, UBE2K, MARCH3, RANGAP1*<br>Immune-associated signaling: *CMIP, SDK1*<br>Transcription regulations: *PMEPA1, BAZ1A, PER2*<br>Signal transduction: *PDE2A, GNA11, NCS1, MOB3B*<br>Solute transport: *SLCO4A1, SLC35E1*<br>Cytoskeleton: *PHLDB1, FNBP1, SIPA1L1,*<br>lncRNA: *AC126696, AC007619* | ↑ FEV1 | **Coordinated homeostatic program integrating signaling, structural organization, proteostatis and transcriptional regulation**<br><br>Not disease-specific, but lung function-preserving |
| **B3** | Transcriptional regulation: *TCF7L2, TLE3, ZNF131, GPBP1L1*<br>Protein turnover and trafficking: *RNF10, AREL1, NUP58, CEP83, ASTL*<br>Post-transcription regulation: *SUGP1, PRRC2C, RC3H1*<br>Mitochondrial function: *GFPT1, SLC25A26, DNM1L, ENOSF1, AGPAT4*<br>Signal transduction: *PHLPP2*<br>DNA repair: *POLH*<br>Cellular structure: *CCDC138* | ↑ FEV1 | Regulation of metabolic signaling pathways, metabolic activity, transcriptional and post-transcriptional governance, genome maintenance mechanisms |
| **B4** | Transcriptional regulators and chromatin control: *EGR1, SNAI1, BRD2, CHD4, HEXIM1, NRARP*<br>Non-proliferation: *CDKN1C* | ↑ Sputum Eosinophils<br>↑ Blood Eosinophils<br>↑ Periostin | Transcription plasticity, cellular state protection, extracellular matrix turnover, restrained proliferation (reduced adipose expansion) |

| | | | |
|---|---|---|---|
| | Signal transduction and second messenger: *PRKCG, AC008397*<br>Oxidative stress protection and protein protection: *AL691403, HSPA2, YME1L1, TRIM69*<br>Extracellular matrix remodeling: *ADAMTS1*<br>RNA modification and regulation: *NSUN6, EMC1-AS1*<br>lncRNA: *KCNQ1OT1, C6orf99, AC004832*<br>Cellular structure: *LRRC23* | ↓ Total Fat Mass | **Not disease specific but associated with T2 inflammation** |
| **B5** | Transcription regulation and chromatin regulation: *INTS6, NFATC3, GATAD2A, GSE1, NFAT5, ASXL1, BAZ2A, BRMS1L, PHF21A*<br>Cell-cycle control: *LIN52, TP53*<br>RNA regulation and post-transcription: *NOCT, AC092849, AL122035, KRTAP5-AS1, AC005480, AC012020* (lncRNA)<br>Cell-cell interactions and cell structure: *EFNB2, EPB41L5*<br>Poorly characterized: *CYB5D1* | ↑ Sputum Eosinophils<br>↑ Blood Eosinophils<br>↑ Periostin<br><br>↓ ICS dose<br>↓ Asthma duration | Genomic stability, transcriptional restraint, and controlled responsiveness over proliferation<br><br>**Maintenance of tissue integrity** |
| **B6** | Energy metabolism: *PYGM*<br>Transcription regulation: *POLR2A, KDM6B, PBRM1, RLF, RFX1, MEF2D, AHDC1*<br>RNA regulation: *NEAT1, LINC01619, HCG18, KIF9-AS1, AL021707*<br>Signal transduction: *GIPR, RAPGEFL1* | ↑ Periostin<br><br>↓ ICS dose | Transcriptional plasticity, chromatin remodeling, and RNA regulation<br><br>**Cellular adaptive state** |

| | | | |
|---|---|---|---|
| | Cell homeostasis and cellular structure: *SLC9A1, SPTAN1, SEC24A*<br>Stress response: *ABTB2*<br>lncRNA: *C3orf20* | | |
| **C7** | Transcriptional regulation: *RFX3, SCMH1*<br>RNA processing and post-transcription: *MBNL1, CELF2, ZCCHC7*<br>Signal transduction: *CASK, INPP4B*<br>Protein turnover and trafficking: *WWP1, VPS41*<br>DNA repair: *ERCC6L2, KIAA1328*<br>Transport: *SLC9A9, FUT8*<br>Immune modulation: *TAFA1, RPS6KA5*<br>lncRNA: *PCAT1, LINC02328*<br>Cell structure: *CEP85L* | ↑ FEV1<br>↑ Sputum Macrophages<br><br>↓ **Asthma diagnosis**<br>↓ Asthma exacerbation<br>↓ Age at asthma onset<br>↓ Asthma duration<br>↓ IgE<br>↓ ΔFEV1 | Gene-expression regulation through transcriptional and RNA-processing mechanisms<br><br>Signal integration, protein trafficking, and genome maintenance<br><br>Immune-regulatory state<br><br>**Cellular stability and immune regulation** |
| **C8** | Gene expression, RNA, transcription: *PAN2, EEFSEC, THOC2, PSMA3-AS1*<br>Genome integrity and DNA: *WRN*, *TOP2B, C15orf41, PPP4R3A*<br>Signal transduction: *RASA2, RAB28*<br>Membrane/cell organization and ion transport/homeostasis: *LRRC8D, TMEM265, TMEM38B, EFR3A*<br>Protein turnover and trafficking: *UBAP1L, NEU3, ARMCX4*<br>Cell integrity: *CEP120, MLLT6*<br>Immune modulation: *DOCK8* | ↑ FEV1<br>↑ Sputum Macrophages<br>↑ BRI<br>↑ W/H<br><br>↓ **Asthma diagnosis**<br>↓ Asthma exacerbation<br>↓ ICS dose<br>↓ age at asthma onset<br>↓ ΔFEV1 | Cellular homeostasis and genome-stabilizing program integrating RNA processing, DNA integrity, controlled signal transduction, and cell organization.<br><br>**Cell-intrinsic maintenance program under stress** |

| | | | |
|---|---|---|---|
| **C9** | <u>Transcription, chromatin remodeling, nucleus</u>: *KAT6A, ARID1B, MED13L, RUNX1, HIVEP2, STAG1*<br><u>RNA processing and transcription</u>: *PAN3, CDK13, CDKAL1*<br><u>Signal transduction/second messenger</u>: *PDE8A, STIM1, TNIK*<br><u>Cell organization, migration, trafficking</u>: *AAK1, BCAS3, ANKRD17, DOCK9, ANKH*<br><u>Protein turnover, apoptosis, stress</u>: *BIRC6, COP1, BTBD9* | ↑ FEV1<br>↑ Sputum Macrophages<br><br>↓ **Asthma diagnosis**<br>↓ Asthma exacerbation<br>↓ ICS dose<br>↓ ΔFEV1<br>↓ asthma duration | Chromatin remodeling, RNA processing, and signal transduction<br><br>Regulation of protein turnover, apoptosis resistance, and cytoskeletal organization<br><br>**Stable and adaptable cellular state** |
| **D10** | <u>Immune signaling/antiviral innate immunity</u>: *TRIM25*<br><u>Response to stress</u>: *MAPK8IP3, SHC1*<br><u>Signal transduction</u>: *AGAP4, RGL1*<br><u>Trafficking, endosome:</u> *SNX14, ANKFY1, TM9SF3*<br><u>Transcription and post-transcription</u>: *POLR3B, AGO1, MTA3, LINC02595*<br><u>Protein turnover</u>: *CUL2, ZYG11B, SIL1*<br><u>Mitochondria and metabolic regulation</u>: *ABCB10, PDHX, PEX26, AGPS*<br><u>Regulation of growth</u>: *CREG1* | ↑ FEV1<br>↑ Sputum Macrophages<br><br>↓ **Asthma diagnosis**<br>↓ Age at asthma onset<br>↓ ICS dose<br>↓ Δ FEV1 | Innate immune-linked stress response with coordinated signaling, trafficking, and metabolic regulation<br><br>**Cellular adaptation,**<br>**maintaining cellular homeostasis** |
| **D11** | <u>lncRNA:</u> *LEF1-AS1, CHRM3-AS2, LINC01725, MAP4K3-DT*<br><u>Inflammatory signaling:</u> *EDAR, ALPK1, TCF7*<br><u>DNA repair:</u> *RNF144A* | ↑FEV1<br>↑ Sputum Macrophages<br><br>↓ **Asthma diagnosis**<br>↓ Sputum Eosinophils | Cellular regulatory and signaling program integrating lnc-RNA, transcription, DNA repair, protein turnover and inflammatory signaling, migration, apoptosis |

| | | | |
|---|---|---|---|
| | <u>Protein turnover</u>: *RNF175*<br><u>Wnt-pathway</u>: *DACT1 (TCF7)*<br><u>Transcription regulation</u>: *GLIS3, ZHX3*<br><u>Apoptosis related signaling</u>: *BEX3, APBB1*<br><u>Cell migration</u>: *ADTRP, PTK2, CLUAP1*<br><u>Mitochondria function/cell metabolism</u>: *BCKDHB* | ↓ Asthma exacerbation<br>↓ Age at asthma onset<br>↓ ICS dose<br>↓ Asthma duration<br>↓ Δ FEV1 | **Maintenance of functional cellular stability, integrating stress, and signaling responses.** |
| **D12** | <u>Intracellular transport:</u> *DYNC1LI2, VAPB, CEP68, PHLDB2, PLEKHM2, SLC66A2, EPN2*<br><u>Lipid metabolism and mitochondria function</u>: *ABCD3, ACSL3, SOD2*<br><u>Cell-cell interaction and immune signaling</u>: *TMEM204, CCR7, CD59*<br><u>Transcription regulation</u>: *TLE2, MYBBP1A*<br><u>Protein control and stress adaptation</u>: *FBXO31, DNAJA4*<br><u>Adipose specific signaling</u>: *PGRMC2* | ↑FEV1<br>↑ Sputum Macrophages<br><br>↓ **Asthma diagnosis**<br>↓ Sputum Eosinophils<br>↓ Asthma exacerbation<br>↓ Age at asthma onset<br>↓ ICS dose<br>↓ Asthma duration<br>↓ Δ FEV1 | Integration of intracellular trafficking, lipid and mitochondrial metabolism, proteostasis, and immune–stromal communication<br><br>**Maintenance of cellular function and metabolic activity** |
| **E13** | <u>Mitochondrial signature:</u> *NDUFA12, NDUFB4, FMC1, ATP5PF, MICOS10*<br><u>Ribosome and protein synthesis</u>: *MRPS21, RPS19BP1, RPS27L, SEM1, DDOST*<br><u>Post-translational</u>: *SUMO3*<br><u>Regulation of cell cycle:</u> *ANAPC11* | ↑ **Asthma diagnosis**<br>↑ Blood neutrophils<br>↑ IgE<br><br>↓ FEV1 | Increased mitochondrial energy production, ribosome biogenesis, protein processing, and cytoskeletal remodeling<br><br>**Metabolically stressed and activated cells** |

| | | | |
|---|---|---|---|
| | Structural cell adhesion and migration: *BRK1, PDLIM2, PDAP1*<br>Cholesterol synthesis, Glycosylation: *SCP2, PMM1* | | |
| **E14** | Mitochondrial function: *ATP5F1A, COA4, TXN2, ETFA, ETHE1*<br>Proteostasis and protein quality control: *PSMC1, PSMC3, DNLZ, TSR3, NOL12*<br>Vesicles, organelles, and lysosome: *TRAPPC5, BLOC1S4, LAMTOR2*<br>Innate immune modulation and tissue remodelling: *MMP25, UBL7* | ↑ **Asthma diagnosis**<br>↑ IgE<br>↑ FeNO<br>↑ Exacerbations<br>↑ ICS dose<br>↑ Age at diagnosis<br><br>↓ FEV1<br>↓ Sputum Macrophages | Mitochondrial hyperactivity, oxidative stress, high protein turnover, and altered lysosomal/mTOR signaling<br><br>**Metabolically stressed and activated cells** |
| **E15** | Ribosomal proteins: RPL/RPS family (19 genes)<br>Translation: *NACA* | ↑ **Asthma diagnosis**<br>↑ ICS dose<br><br>↓ FEV1 | Increased ribosome abundance and protein synthesis capacity<br><br>**Metabolically stressed and activated cells** |
| **E16** | Mitochondrial signature: *COX6C, NDUFB8, UQCR11, ATP5F1E, ATP5MG*<br>Protein processing: *PSME1, SPCS1, UBL5*<br>Cytoskeleton and contractility: *ARPC3, TPM3, MYL6*<br>Immune and antigen-presentation markers: *B2M, GZMA, EMP3*<br>Transcriptional, chromatin regulation: *DRAP1, ELOB, MEAF6, H3F3A* | ↑ **Asthma diagnosis**<br>↑ Exacerbations<br>↑ ICS dose<br>↑ ΔFEV1<br>↑ Years with asthma<br><br>↓ FEV1<br>↓ Sputum macrophages | Increased energetic demand, cytoskeletal reorganization, and low-level immune engagement<br><br>Pyroptosis<br><br>**Structural and functional remodeling** |

| | | | |
|---|---|---|---|
| **E17** | Interferon signaling and innate immune activation: *GBP1, PYHIN1, ZC3H11A, GSR*<br>Immune cell signaling and activation: *SLAMF6, SLAMF7, PLEK, MIEN1*<br>Chemokine signaling: *CCL3/MIP-1α*<br>Protein quality control and signaling: *FBXO6, MIB2, PPM1M* | ↑ **Asthma diagnosis**<br>↑ ΔFEV1<br><br>↓ FEV1 | **Interferon signaling, innate immune activation, and chemokine-mediated recruitment** |
| **E18** | T2 immune programming: *GATA3*<br>Intracellular trafficking and secretion: *VAMP4, ERGIC2, DYNLT3*<br>Stress and hypoxia signaling: *SIAH2, CCS, UQCC2*<br>Signal regulation: *PTPN1*<br>Transcriptional and RNA processing: *ZRANB2, DHX30, RPA3, MYCBP* | ↑ **Asthma diagnosis**<br>↑ Exacerbations<br>↑ ICS dose<br>↑ age at diagnosis<br>↑ years with asthma<br>↑ ΔFEV1<br><br>↓ FEV1<br>↓ Sputum Macrophages | **Th2-associated transcriptional programming, altered intracellular trafficking, and stress signaling** |
| **E19** | T-cell activation and stimulation: *CRTAM, TNFRSF9, ELF1*<br>Immune regulation and migration: *RGS1, BTG1*<br>Cell survival, stress and mitochondria: *SELENOK, MANF, GRPEL1*<br>RNA processing and translational control: *SNU13, DDX18, DDX24, RPP21*<br>Autophagy: *WDR45*<br>Proteostasis: *ADRM1* | ↑ **Asthma diagnosis**<br>↑ Exacerbation<br>↑ ICS dose<br>↑ asthma diagnosis<br>↑ age at diagnosis<br>↑ years with asthma<br>↑ ΔFEV1<br><br>↓ FEV1<br>↓ Sputum Macrophages | **T-cell activation and survival programs** |
| **F20** | Immune and inflammatory response: *IL17RC, LY96, HRH2, GP6, GRN* | ↑ Platelet count<br>↑ TG/HDL<br>↑ Blood neutrophils | Immune signaling, vesicle trafficking, metabolic transport, and oxidative stress |

| | | | |
|---|---|---|---|
| | Cellular metabolism and transport: *SLC35F6, SLC16A5, CA13*<br>Cell signaling and membrane: *RAB31, CCDC50, STOML1, TMEM69, LIMS2, NDRG2*<br>Oxidative stress/Ferroptosis: *AIFM2, EARS2*<br>DNA repair: *SLX1A*<br>Proteostasis: *ASB13*<br>lncRNA: *HEIH*<br>Cell structure: *NEB* | ↑ IgE<br>↑ Exacerbations | **Sustained inflammatory activation supported by altered cellular metabolism** |

FEV1 – forced expiratory volume in 1 second. ICS – inhaled corticosteroid. Ig – immunoglobulin. Δ – post-bronchodilator change. BRI – body roundness index. W/H – waist-to-hip ratio. TG/HDL – triglyceride-to-HDL ratio.

Supplemental Table 2:

**Asthma-specific T/NK iterativeWGCNA Module Key Gene Sets**

| **Module** | **Key Genes (20)** | **Clinical correlation** | **Integrated interpretation** |
|---|---|---|---|
| **P-50** | Cell polarity/organization, cytoskeleton: *ANKRD6, PARD3B, NAV2, EPN1, PLEK2*<br>Cellular metabolism: *CPA6, RRAGD*<br>Intracellular transport: *IFT140, CFAP97D2*<br>Transcription regulation: *ZNF117, SPIN3, ZNF440*<br>LncRNA and poorly characterized: *AC048382, AL096803, CBWD3, AC010618, RTL8, AL117336*<br>Adipogenesis: *AAMDC* | ↑ HbA1c<br>↑ Age at asthma diagnosis<br>↑ IL-1β<br><br>↓ ΔFEV1 | Nutrient sensing, ciliary and polarity signaling, remodeling |
| **S-71** | Transcription and post-transcription: *PRPF40B, CNOT10, YY1AP1, EIF3A, SAMD1, MAFG*<br>Cytokine signaling, immune regulation: *SOCS3, IKZF4, GIMAP8*<br>Regulatory and non-coding RNA: *MIR17HG, AC004889, AC138965, AP000829, AC005726, AL365203, SLC25A25-AS1*<br>Cell homeostasis, structure and adhesion: *GJC2, PKD1, BBS1, ITGA* | ↑ Periostin | Integration of transcriptional control, cytokine signaling, RNA regulation, and cell adhesion/ECM interaction.<br>**Tissue remodeling and immune regulation** |
| **S-72** | Extracellular matrix remodeling: *ADAMTS1* | ↑ Periostin<br>↑ Blood eosinophils<br>↑ Sputum eosinophils | Transcription plasticity |

| | | | |
|---|---|---|---|
| | Transcriptional regulators and chromatin control: *BRD2, EGR1, SNAI1, CHD4, HEXIM1, NRARP*<br>lncRNA: *KCNQ1OT1, C5orf17*<br>Signal transduction and second messenger: *PRKCG, AC008397*<br>RNA modification and regulation: *EMC1-AS1, NSUN6*<br>Oxidative stress protection and protein protection: *AL691403, HSPA2, TRIM69*<br>Regulators of cell proliferation: *CDKN1C, PIK3R3*<br>Protein trafficking**:** *ARL17A, RND1* | | Cellular state protection, extracellular matrix turnover, restrained proliferation (reduced adipose expansion) |
| **S-73** | Metabolic stress signaling: *AKT1S1, SGK1, DENND4C*<br>Cell morphology and cytoskeleton: *RHOU, TBC1D31, MYO19, WDR5B*<br>Transcription regulation: *KDM3B, ZFHX3*<br>Extracellular matrix and tissue structure: *PLOD1, LGALS4*<br>Poorly characterized elements: *RTL5, SPATS2, CYB5D2, ANKRD33B, AC109927, AC004083*<br>lncRNA: *THSD4-AS1, DHRS4-AS1*<br>Copper transport: *SLC31A1* | ↑ Sputum eosinophils, Black, Hispanic<br><br>↓ White | Coordinated stress and remodeling response |
| **V-81** | Transcription regulations: *KLF7, CREG1, AKNAD1, ZCCHC24, POLR1B, PLAGL1* | ↑ IgE | Transcriptional control, regulated signaling, and extracellular matrix turnover |

| | | | |
|---|---|---|---|
| | Signal transduction/ signaling: *RGL1, SGIP1, EFCAB13, ASB3, PACS2, AGAP4*<br>LncRNA and poorly characterized: *LINC02595, AC006141, SEMA6A-AS1, AC100793*<br>ECM remodeling and proteolysis: *TIMP2, CTSB, CTSL*<br>Cell adhesion: *PPFIA1* | | |
| **V-83** | Cellular stress, survival: *AIFM2, CRYAB, MTRNR2L8, NDRG2, SLX1A*<br><br>Cell signaling and membrane: *LIMS2, RAB31, CCDC50, MYRIP, SOWAHC*<br>Immune and inflammatory response: *HRH2, GRN*<br>Metabolic transport and lipid signaling: *FFAR3, SLC16A5, SLC35F6*<br>Epigenetic and lncRNA: *LINC01376, HEIH, MIR4458HG, FP236383, COPRS* | ↑ IgE<br>↑ TG/HDL<br>↑ Blood neutrophils<br>↑ Platelets<br>↑ hsCRP<br>↑ Exacerbations | Stress-adapted inflammatory program that integrates metabolic dysregulation with innate immune signaling<br><br>Stress responses, lipid and metabolite sensing |
| **V-84** | Cellular metabolism and membrane biosynthesis: *ETNK1, MTRR, FKTN*<br>RNA processing and transcriptional/post-transcriptional regulation: *NOP9, ZNF443, ZRANB2-AS2, AC002383, AL096870, AL117336, DHX32*<br>Cytoskeletal organization and vascular-associated signaling: *ASTN2, PRKG1, EVI5* | ↑ TG/HDL<br>↑ Platelets | Membrane remodeling, transcriptional stabilization, and cytoskeletal organization |

| | |
|---|---|
| | Extracellular matrix and stromal–vascular interface: *ADAMTSL5*<br>Cellular signaling and protein turnover: *SLC35G1, STK32C, NAV1, FBXL7*<br>Innate immune signaling: *TRIM25* |
| FEV1 – forced expiratory volume in 1 second. Ig – immunoglobulin. Δ – post-bronchodilator change. TG/HDL – triglyceride-to-HDL ratio. | |

Supplemental Table 3:

**Myeloid iterativeWGCNA Metamodule Key Gene Sets**

| **Metamodule** | **Key genes** | **Clinical correlation** | **Integrated Interpretation** |
|---|---|---|---|
| **AA** | lncRNAs and uncharacterized transcripts (regulatory): *LINC01366, LINC01033, AC016134, AC008397, AC114977, AC136604 , AC080013, AL139106, AL049869, AL591518, AL035458, AP000640, DNAJB5-DT*<br>Ciliary structure and motility: *DNAI2, SAXO1, LRRC23*<br>Vesicle trafficking and secretion: *RAB26, PARP16*<br>Cytoskeleton and structural organization: *PRX, C1orf189* | ↑ Sputum Eosinophils, Blood Eosinophils, periostin<br><br>↓ Asthma diagnosis, severity and duration (AA-3 module only) | Regulatory/adaptive tissue interface module |
| **BB** | Intracellular signaling (immune activation and survival): *DENND5A, MAP2K1, GAB2, NEDD9* | ↑ Fasting glucose, VAT mass, FEV1% predicted, Sputum macrophages<br><br>↓ Asthma diagnosis, asthma severity, age at | Macrophage-driven immunoregulatory and metabolic adaptation module, potentially reflecting a compensatory systemic response despite metabolic activation |

| | | | |
|---|---|---|---|
| | <u>Transcriptional and epigenetic regulation:</u> *CREBBP, CUX1, ELF4, MXD1, MRTFA*<br><u>Cytoskeleton remodeling and cell migration:</u> *FMNL1, DIAPH1, PAFAH1B1*<br><u>Vesicle trafficking and RNA/post-transcriptional control:</u> *AGO2, AGO3, LARP4B*<br><u>Immune modulation and extracellular interaction:</u> *CHST15, CMIP, BTBD9, ANKRD17* | asthma onset, asthma duration, ΔFEV1, asthma exacerbations, sputum eosinophils<br><br>↑ IL-6 (BB-13) | |
| **CC** | <u>Cytosolic ribosome and protein translation machinery:</u> *RPS15A, RPL19, RPL32, RPL24, RPL11, RPL18, RPL35A, RPLP2, RPL6, RPS15, RPS24, RPS13* [Components of the large and small ribosomal subunits. Drive mRNA translation → protein synthesis. Often tightly co-expressed in highly anabolic or stressed cells]<br><u>Mitochondrial oxidative phosphorylation (OXPHOS):</u> *NDUFA13, NDUFA11, COX7A2, COX8A, ATP5PB, CHCHD2*<br><u>Protein turnover:</u> *PSMB6*<br><u>Protein modification:</u> *UFC1* | ↑ Asthma diagnosis, disease severity, duration<br><br>↓ VAT mass, FEV1% predicted, Sputum macrophages, sputum and blood eosinophils | Pro-inflammatory, metabolically hyperactive macrophage state linked to asthma severity and tissue dysfunction |
| **DD** | <u>Monocyte / macrophage / dendritic cell lineage (Core innate immune signature):</u> *C1QC, CD209, CD163L1, CMKLR1, CHID1, CTSF, DAB2, SLC40A1* | ↑ Asthma diagnosis, severity, age at asthma diagnosis, ΔFEV1, asthma exacerbations, blood neutrophils, | Pathogenic innate immune–remodeling module tightly coupled to type 2 high asthma severity |

| | | | |
|---|---|---|---|
| | Type 2 inflammation context: *MAF, CD28, CHID1*<br>Tissue remodeling: *LTBP2, HTRA1, PLXND1*<br>Vesicle trafficking: *SCAMP2*<br>Mitochondrial and cellular stress (hypoxia response): *FUNDC1*<br>Signaling / regulatory: *ARHGAP18, PRKACB*<br>Miscellaneous: *SCN9A, LGI2, TMEM106B* | sputum and blood eosinophils, periostin, IgE, FeNO<br><br>↓ VAT mass, fasting glucose, FEV1% predicted, Sputum macrophages | |
| **EE** | Lipid metabolism and cholesterol handling: *LPL, NPC2*<br>Mitochondrial energy metabolism and oxidative capacity: *MRPL15, NDUFB6, GCSH, MDH1*<br>ER protein processing, glycosylation and trafficking: *DPM2, DAD1, GPAA1, PDIA4, TMED9*<br>Immune activation, macrophage activity and stromal interaction: *CD68, MYDGF, MME, ITGA3*<br>Redox balance, heme metabolism and metabolic cofactors: *BLVRB, PCBD1, NENF*<br>Transcriptional regulation and apoptosis control: *SPOCD1, TFPT* | ↑ Asthma diagnosis, ICS dose, age at asthma diagnosis, ΔFEV1, exacerbations, asthma duration<br>↑ FMI and White (EE-31)<br><br>↓ Sputum macrophages, FEV1, VAT mass, fasting glucose | Lipid-driven macrophage metabolic stress module that integrates adipose dysfunction with asthma severity |

FEV1 – forced expiratory volume in 1 second. VAT – visceral adipose tissue. Ig – immunoglobulin. Δ – post-bronchodilator change. FeNO – fractional exhaled nitric oxide. ICS – inhaled corticosteroid. TG/HDL – triglyceride-to-HDL ratio.
