## Supplemental Figure 1 for "Adipose tissue as a site of immune activation and dysfunction in individuals with obesity and asthma"

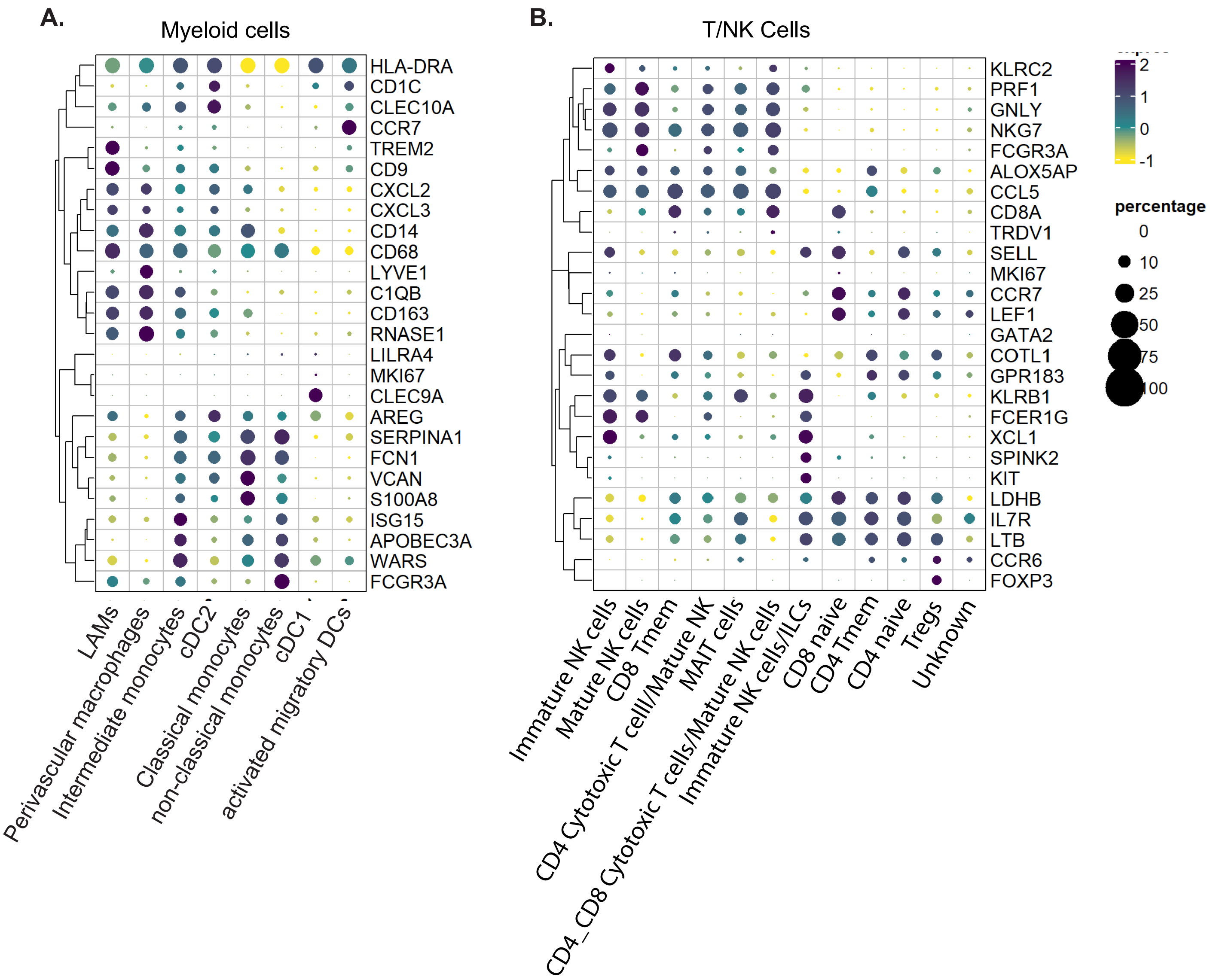


**Supplemental Figure 1** (related to all Figures) – Key genes used to annotate cell type for myeloid cells (panel A) and T/NK cells (panel B).
