## Supplemental Figure 2 for "Adipose tissue as a site of immune activation and dysfunction in individuals with obesity and asthma"

A.

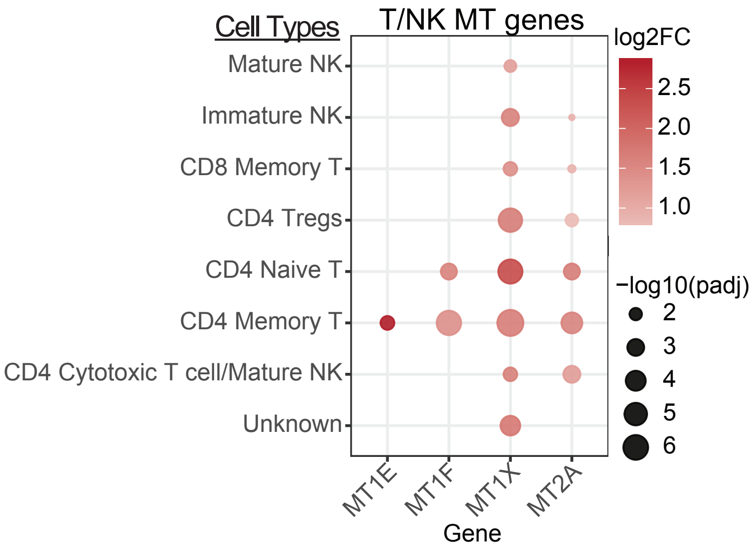


B.

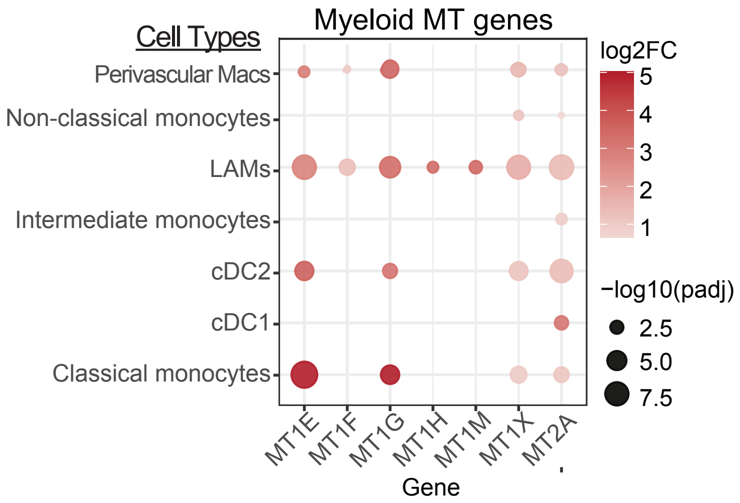


**Supplemental Figure 2** (related to Figures 2-3)- Effector cell signature supports dysfunctional phenotype. (A) Pseudo-bulk metallothionine gene expression in T/NK cluster by cell subset with non-asthma set as reference group. (B) Pseudo-bulk metallothionine gene expression in myeloid cluster by cell subset with non-asthma set as reference group.
