## Supplemental Figure 3 for "Adipose tissue as a site of immune activation and dysfunction in individuals with obesity and asthma"

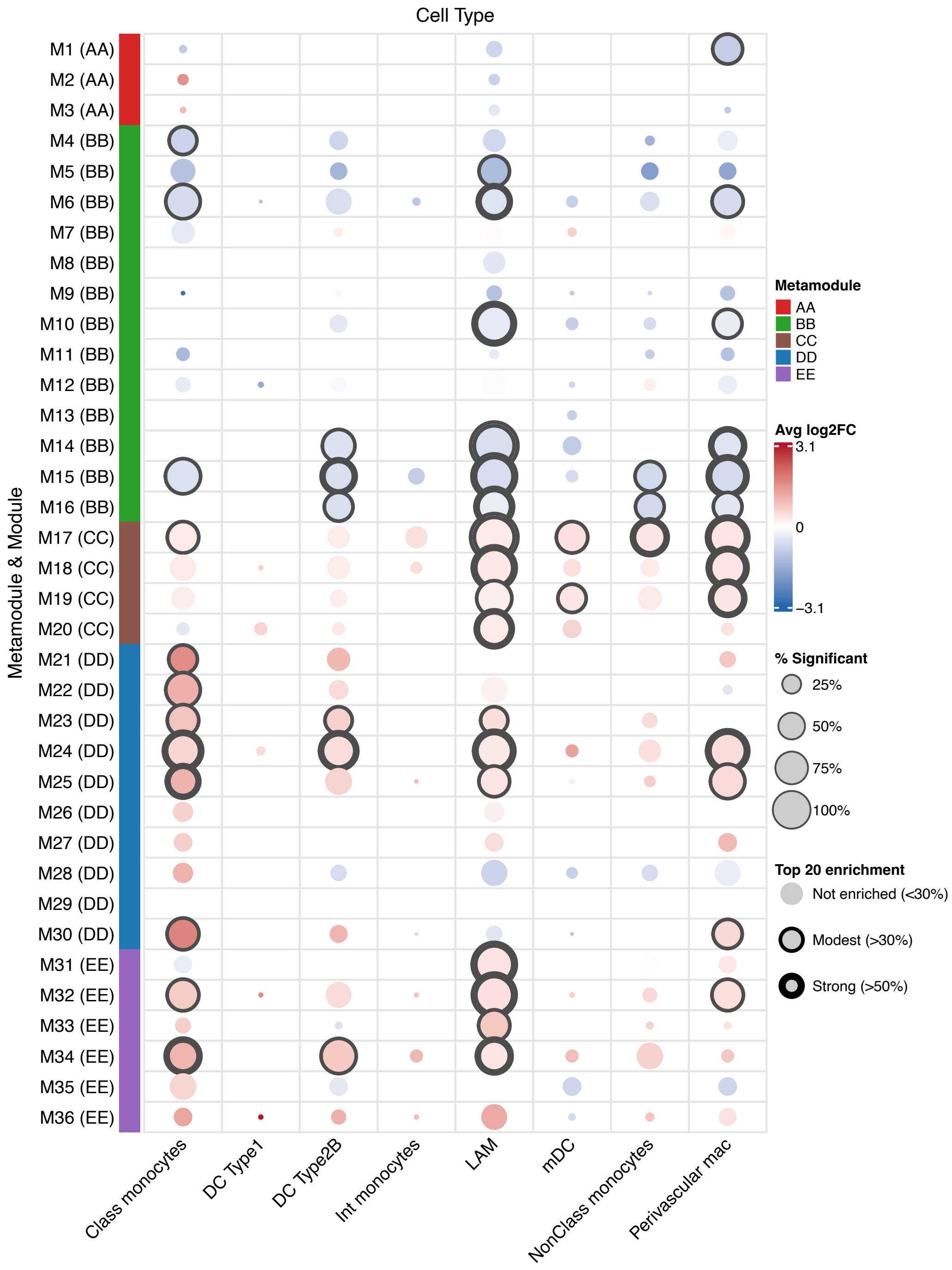


**Supplemental Figure 3 (related to Figure 7) – Genes from specific cell-types for metamodules.** Integration of cell-type specific pseudobulk DE data for each metamodule using kME threshold ≥0.75 with non-asthma set as reference. Percent of genes from each cell-type meeting threshold for adjusted p-value significance and represented in the top 20 enriched genes in each metamodule denoted.
